# Spatial characterization of interface dermatitis in cutaneous lupus reveals novel chemokine ligand-receptor pairs that drive disease

**DOI:** 10.1101/2024.01.05.574422

**Authors:** Saeed Shakiba, Nazgol-Sadat Haddadi, Khashayar Afshari, Janet E. Lubov, Haya S. Raef, Robert Li, Ümmügülsüm Yildiz-Altay, Mridushi Daga, Maggi Ahmed Refat, Evangeline Kim, Johanna Galindo de Laflin, Andressa Akabane, Priscilla Romano, Jane Vongvirath, Shany Sherman, Elizabeth MacDonald, James P. Strassner, Liang Zhang, Michael Leon, Christina E. Baer, Karen Dresser, Yan Liang, James B Whitley, Sladjana Skopelja-Gardner, John E Harris, April Deng, Matthew D. Vesely, Mehdi Rashighi, Jillian Richmond

## Abstract

**Background:** Chemokines play critical roles in the recruitment and activation of immune cells in both homeostatic and pathologic conditions. Here, we examined chemokine ligand-receptor pairs to better understand the immunopathogenesis of cutaneous lupus erythematosus (CLE), a complex autoimmune connective tissue disorder.

**Objectives:** Our objectives were to (1) characterize the cellular and proteomic constitution of interface dermatitis in CLE using blister biopsies, (2) map chemokine:ligand receptor pairs that govern recruitment of immune cells to form interface dermatitis in CLE, and (3) perform unbiased analyses in tandem on different clinical subtypes to identify novel genes and proteins underlying discoid versus subacute CLE.

**Methods:** We used suction blister biopsies to measure cellular infiltrates with spectral flow cytometry in the interface dermatitis reaction, as well as 184 protein analytes in interstitial skin fluid using 96-plex immunoassay targeted proteomics. Flow and 96-plex immunoassay data concordantly demonstrated significant increases in T cells and antigen presenting cells (APCs). We also performed spatial transcriptomics and spatial proteomics of punch biopsies using digital spatial profiling (DSP) technology on CLE skin and healthy margin controls to examine discreet locations within the tissue.

**Results:** Spatial and 96-plex immunoassay data confirmed elevation of interferon (IFN) and IFN-inducible CXCR3 chemokine ligands. Comparing involved versus uninvolved keratinocytes in CLE samples revealed upregulation of essential inflammatory response genes in areas near interface dermatitis, including *AIM2*. 96-plex immunoassay data confirmed upregulation of Caspase 8, IL-18 which is the final product of AIM2 activation, and induced chemokines including CCL8 and CXCL6 in CLE lesional samples. Chemotaxis assays using PBMCs from healthy and CLE donors revealed that T cells are equally poised to respond to CXCR3 ligands, whereas CD14+CD16+ APC populations are more sensitive to CXCL6 via CXCR1 and CD14+ are more sensitive to CCL8 via CCR2.

**Conclusions:** Taken together, our data map a pathway from keratinocyte injury to lymphocyte recruitment in CLE via AIM2-Casp8-IL-18-CXCL6/CXCR1 and CCL8/CCR2, and IFNG/IFNL1-CXCL9/CXCL11-CXCR3, and identify potential novel biomarkers of disease.

**Graphical abstract:** 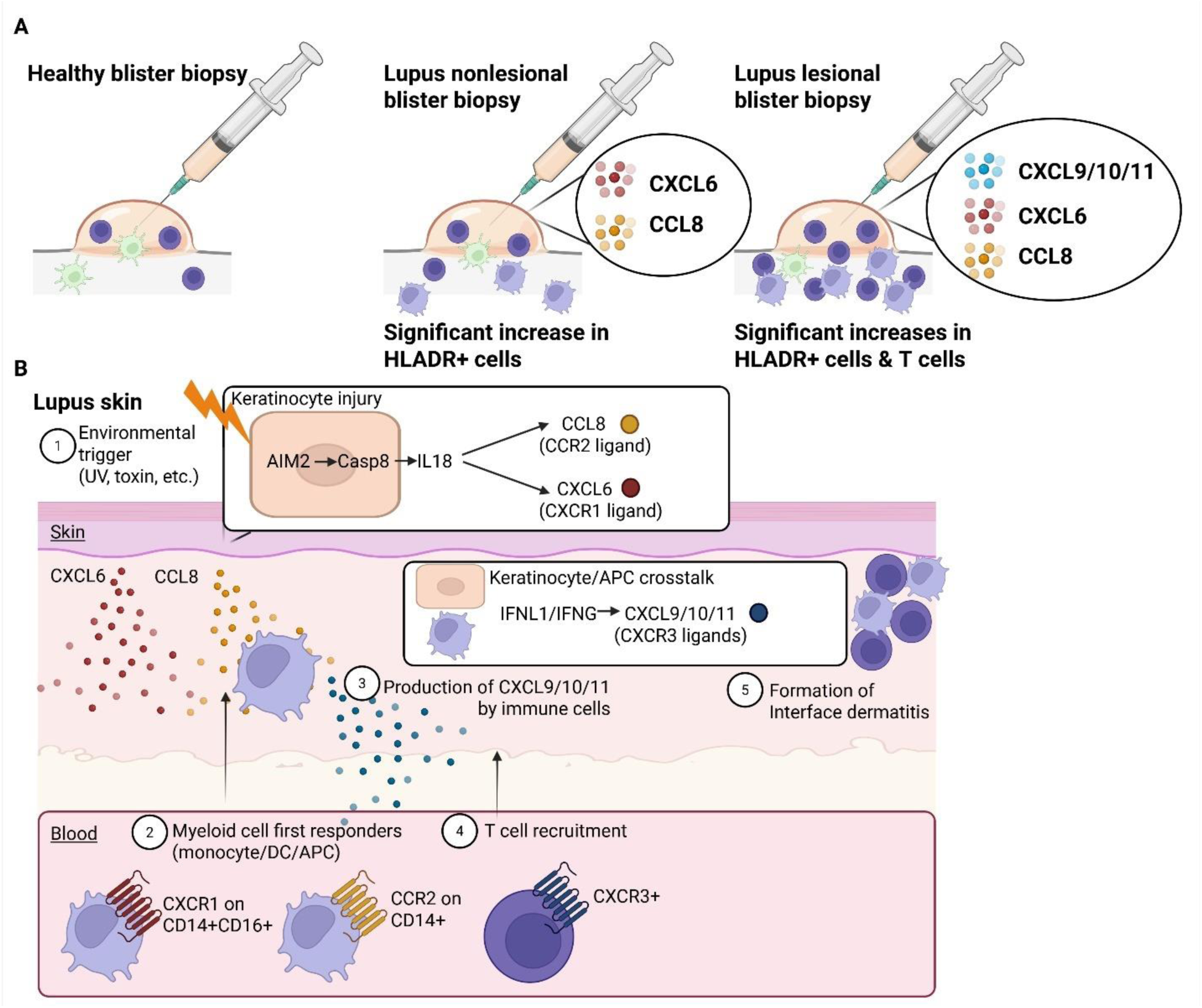

**Model of chemokine systems governing recruitment of immune cell subsets to form interface dermatitis in cutaneous lupus**. A. Summary of fresh tissue blister biopsy studies demonstrating increases in HLADR+ cells in nonlesional lupus biopsies as assessed by flow cytometry, and increased CXCL6 and CCL8 as assessed by 96 plex immunoassay. Lesional lupus biopsies also demonstrated significant increases in T cells and CXCL9/10/11 production. B. Model of chemokine-directed formation of interface dermatitis in cutaneous lupus. 1. Whole transcriptome atlas (WTA) digital spatial profiling (DSP) revealed increased *AIM2* in keratinocytes proximal to inflammation, which is reported to be induced by environmental triggers including UV light and toxins. We also noted increased Caspase 8 (Casp8) and IL18 at the protein level, which can be induced downstream of AIM2. Chemokines including CXCL6 and CCL8 can be induced downstream of IL18, explaining how CCL8 and CXCL6 might be induced. 2. Recruitment of myeloid cell first responders by CCL8 and CXCL6. CD14+CD16+ myeloid cells, which were recently described in nonlesional lupus skin, express CXCR1 and migrate towards CXCL6, whereas CD14+CD16-myeloid cells express more CCR2 and migrate towards CCL8. 3. The CXCR3 ligands CXCL9/10/11 are expressed by keratinocytes, but more strongly in CD45+ immune cell and T cell regions of interest (ROIs). 4. T cells express CXCR3 and migrate towards CXCL9 and to a greater extent CXCL11. 5. The recruited HLADR+ myeloid populations and T cells contribute to formation of interface dermatitis. Thus, we propose a model in which keratinocyte/myeloid crosstalk can reinforce chemokine systems to optimally recruit lymphocytes and other immune cells to form interface dermatitis. Created with Biorender.com.

**Plain language summary:** Lupus skin rashes arise during flares, after exposure to medications or sunlight, or in response to other triggers of inflammation. To understand how white blood cells enter the skin to cause these rashes, we used new technologies to look at proteins that attract them into the skin. We found proteins that are expressed by skin cells in lesions that can recruit specific types of white blood cells that are thought to be the initiators of skin rashes. Once in the skin, these and other white blood cells can make additional proteins that bring in more and more cells. We hope that our findings will be used to test new topical treatments for lupus and other autoimmune skin rashes.

## INTRODUCTION

Cutaneous lupus erythematosus (CLE) is a spectrum of clinically diverse autoimmune connective tissue disorders with shared histopathological features including interface dermatitis and lupus band reaction^1^. Chronic cutaneous lupus erythematosus (CCLE, most common sub-entity discoid lupus erythematosus (DLE)), subacute cutaneous lupus (SCLE), and acute cutaneous lupus erythematosus (ACLE) include the majority of cutaneous lupus clinical subtypes, though there are other more rare entities which all have varying degrees of association with systemic lupus erythematosus (SLE). In clinical practice, CLE can be refractory to SLE treatments^2–4^, but successful treatment of cutaneous disease may significantly decrease the risk of systemic involvement^5^.

A common feature of CLE is the presence of interface dermatitis, which refers to inflammation and/or degenerative changes at the dermal-epidermal junction. The mechanisms by which immune cells are recruited to form interface dermatitis in CLE are unknown. Previous studies have found that interferon (IFN)-inducible chemokines are highly upregulated in CLE lesions ^6,7^. One recent study characterized the associations of specific chemokine receptors and ligands in T cell subsets in cutaneous versus systemic lupus, demonstrating different profiles in SLE, DLE and SCLE^8^. The cellular sources of chemokine ligands binding to cognate receptors have not been fully characterized.

To date, many CLE studies have focused on transcriptomics including bulk RNA sequencing and single cell RNA sequencing^9^, but overlooked proteomics approaches and cellular analyses. Here, we used a blister biopsy technique^10^ to sample interstitial skin fluid and cells from the interface dermatitis reaction from CLE patients and compared these to healthy donors. Blister fluid does not require enzymatic digestion for analysis, yielding high-quality liquid and cellular biopsies. We used 96-plex immunoassay targeted proteomics to assess inflammatory biomarkers in interstitial skin fluid, and flow cytometry to assess infiltrating immune cells and their phenotypes. We demonstrate significant increases in T cells and HLA-DR+ cells in blister fluid from patients compared to healthy controls. To understand how these immune cells might enter the skin, we assessed chemokine ligand and receptor expression patterns. We confirmed upregulation of CXCL9 and CXCL11 in patient blister fluid and skin biopsies, with expression of the cognate receptor CXCR3 on T cells on both healthy and lupus blood donors. T cells from healthy and lupus patients migrated towards CXCL9 and CXCL11 to varying degrees. We also identified novel chemokine ligands that were elevated in lesional versus nonlesional lupus blister fluid, namely CCL8 and CXCL6. CCL8 binds to the receptors CCR1, CCR2, CCR3 and CCR5. We found that CD14+ monocytes expressed higher CCR2 than other populations of myeloid cells and migrated towards CCL8. In contrast, CD14+CD16+ myeloid cells expressed the CXCL6 receptor CXCR1 and migrated towards this ligand. We also performed matched spatial transcriptomics and proteomics of DLE and SCLE archival biopsies to identify the general cellular sources (keratinocytes vs immune cells) and locations of chemokines and other immune proteins. We identified *AIM2* and *IFNG* in keratinocytes, which can lead to downstream production of CXCL6, CCL8, and CXCL9/11 respectively. Notably, these ligands are not expressed in healthy skin, meaning that immune cells are poised to respond to these ligands in response to injury or insult such as UV light exposure. Further, spatial data identified unique gene and protein expression signatures which may underlie differences in the discoid and subacute clinical subtypes of CLE, in addition to unique expression of B7-H3 protein on SCLE T cells. Importantly, we identify potential treatment targets in both immune cells and keratinocytes that may be of use for future clinical trials, namely *IRAK*s and *CAPN*s and *TYK2*, as well as a novel functional target CXCR1 which we demonstrate can be inhibited with reparixin.

## MATERIALS AND METHODS

### Study subjects

This study was performed with internal review board (IRB) approved protocols at the UMass Chan Medical School (H-14848 and H00021295 for fresh blood and skin tissue; H00020503 for archival skin tissue), Yale University Institutional Review Board (Human Investigative Committee no. 15010105235 for archival skin tissue), and/or Dartmouth Hitchcock Medical Center (STUDY02001542 for PBMCs), and all samples were obtained with written informed consent and were de-identified before use in experiments. Inclusion/exclusion criteria were as follows: subjects with a diagnosis of cutaneous lupus erythematosus by clinical exam performed by a dermatologist were included. Subjects with recent onset of new lesions and objective clinical signs of activity, specifically erythema and/or hair loss/alopecia, were recruited to represent active disease. Patients on treatment were included in the study, as most lupus patients must remain on maintenance therapy.

### Statistics

Statistical analyses were performed using Prism software version 9 (GraphPad) for flow cytometry, NPX software for 96-plex immunoassay, and GeoMX software for spatial datasets. For comparisons across disease states and cell types, two-way ANOVAs with main row effects comparisons were performed to identify highly significant biomarkers or cell types for further analysis. Differences were considered significant at a P value of less than 0.05. DSP statistical analyses using Linear Mixed Model (LMM) or unpaired T-tests were performed in GeoMX software with the resulting p-values that were adjusted for multiple comparisons with the Benjamini-Hochberg procedure (FDR, false discovery rate) and considering the size of regions of interest (ROIs) and slide scanning batch effect. Box plots and volcano plots were generated in GraphPad Prism version 9. P-value or FDR <0.05 were considered significant, with P<0.01 as highly significant. We also used one-way ANOVA or two-way ANOVA for some comparisons in a group of genes between healthy and disease ROIs to answer specific hypotheses generated by -omics datasets.

## RESULTS

### Integrating spatial and -omics approaches for characterizing the inflammatory infiltrate in cutaneous lupus interface dermatitis

We combined spatial studies of archival tissue with analyses of fresh tissue biopsies to validate chemokine and other targets at the protein level (workflow **Fig 1A**). Blister biopsies were used to sample interstitial skin fluid and cells from CLE patients and healthy donors (**Fig 1B, Table S1**). 96-plex immunoassay targeted proteomics on the interstitial skin fluid confirmed previously reported increased levels of IFNg, CXCL9 and CXCL11 in CLE skin biopsies (**Fig 1C**) ^6,8^. 96-plex immunoassay also identified higher levels of CXCL6 and hepatocyte growth factor (HGF), which can increase CXCR3 expression on T cells ^11^, in CLE fluid compared to healthy controls, previously not reported in CLE skin analyses. A 19 color flow panel was used to identify cell types of interest by both UMAP and manual gating. Spectral flow cytometry analysis of blister fluid cells revealed a predominantly lymphocytic infiltrate, with increases in myeloid populations as compared to healthy (**Fig 1D**).

**Fig. 1.**
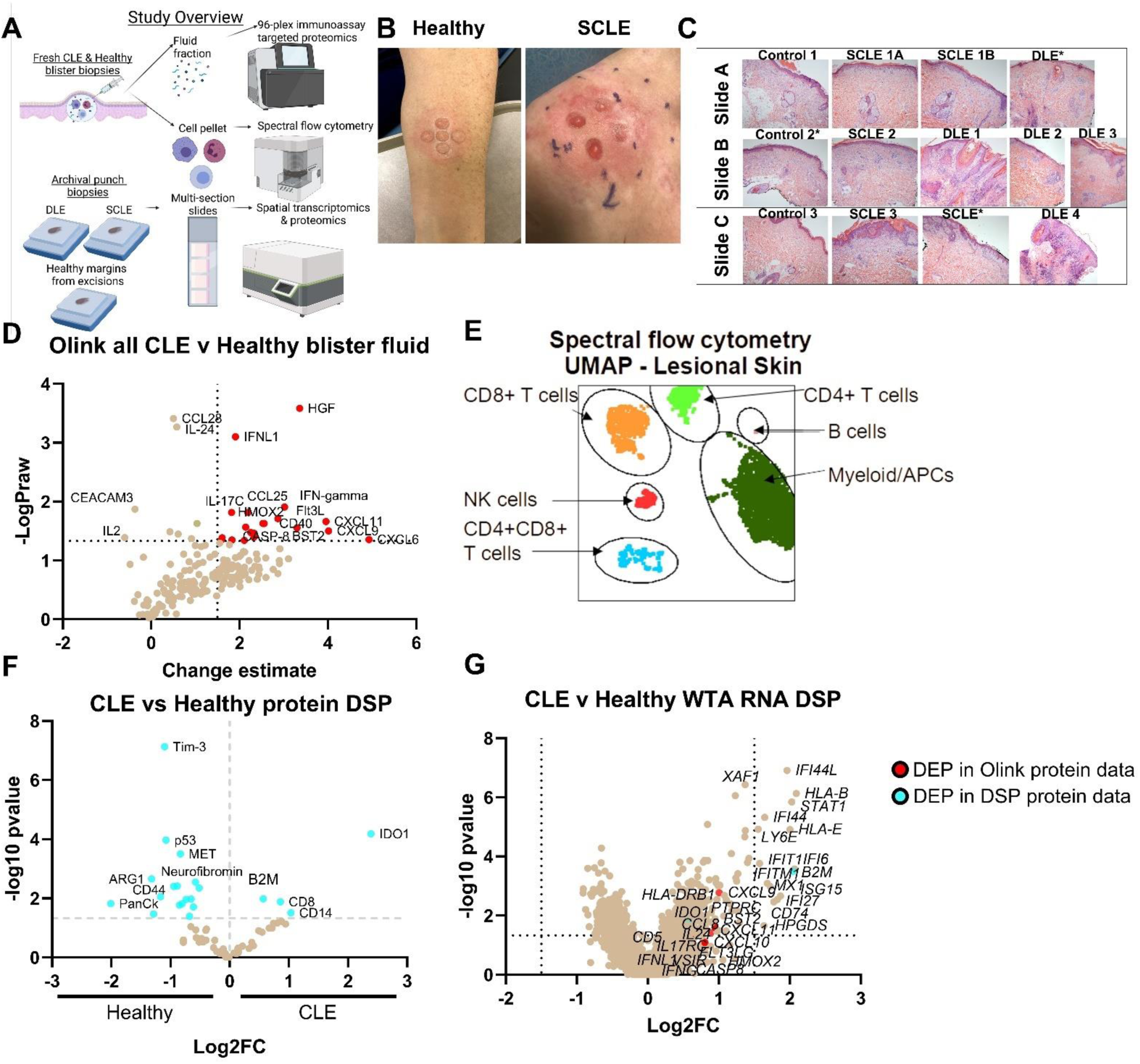
Integration of high-resolution immune techniques with spatial approaches for studying cutaneous lupus erythematosus. (**A**) Study overview demonstrating workflow. We performed blister biopsies on cutaneous lupus erythematosus patients and healthy donors and analyzed the interstitial skin fluid using 96-plex immunoassay proteomics and the cell pellets using Aurora spectral flow cytometry. We also employed biobanked FFPE skin biopsies from DLE, SCLE and healthy margins for NanoString digital spatial profiling (DSP) for protein using 6 modules, and RNA using the whole transcriptome atlas (WTA). Created with Biorender.com. (**B**) Sample photographs of blister biopsies (n=3 healthy and 4 CLE patients). (**C**) H&E images of the slide set used for spatial transcriptomics and proteomics. (n=3 healthy margin controls, 5 SCLE biopsies from 4 patients, and 4 DLE biopsies from 4 patients. * indicates tissue was not fully seated in imaging area for WTA run). (**D**) Volcano plot of 96-plex immunoassay data from the inflammation and neuroexploratory panels of all CLE blisters versus healthy controls (n=7 CLE lesional, 7 CLE nonlesional blisters from 4 donors, and 3 healthy blisters). (**E**) Sample UMAP of immune infiltrates in CLE lesional skin. (representative patient of 4 CLE patients) (**F**) Volcano plot of DSP protein data for all CLE regions of interest (ROIs) versus healthy margin ROIs. (n=95 ROIs assessed from 4 DLE, 3 SCLE and 3 healthy margin controls for protein DSP) (**G**) Volcano plot of DSP RNA WTA data for all CLE ROIs versus healthy margin ROIs. Red dots are also differentially expressed proteins (DEPs) in the 96-plex immunoassay protein dataset, and cyan dots are also DEPs in the DSP protein dataset. (n=41 ROIs assessed from 4 DLE, 3 SCLE, and 2 healthy margin control biopsies for WTA DSP.) (FFPE, formalin fixed paraffin embedded; DLE, discoid lupus erythematosus; SCLE, subacute cutaneous lupus erythematosus; WTA, whole transcriptome atlas; DSP, digital spatial profiling; UMAP, Uniform Manifold Approximation and Projection; DEP, differentially expressed protein; DEG, differentially expressed gene; ROI, region of interest).

Spatial transcriptomics and proteomics on archival tissue biopsies was performed using NanoString Digital Spatial Profiling (DSP, **Fig 1E, Table S2**). Spatial proteomics captured enrichment of CD8 and CD14 as well as beta-2 microglobulin (B2M) and indoleamine 2,3-dioxygenase (IDO) (**Fig 1F**, dataset QC in **Fig S1,** QC of regions of interest (ROIs) in **Fig S2**). Spatial transcriptomics analysis using the Whole Transcriptome Atlas (WTA) of aggregate ROIs recapitulated IFN signatures, and also harmonized with protein biomarkers identified in 96-plex immunoassay and protein DSP (**Fig 1G**). We also validated our DSP WTA dataset using a Cancer Transcriptome Atlas (CTA) dataset (**Table S3**), as well as a bulk RNA microarray dataset previously generated from matched tissue blocks (**Fig S3 & 4**). These datasets had many shared differentially expressed genes (DEGs), including *B2m, Cxcl9, Cxcl10* and other IFN- and immune-related genes.

### Characterization of the immune infiltrate in CLE interface dermatitis

We sought to characterize which cells comprised the interface dermatitis in CLE using multimodal approaches. To do this, we performed spectral flow cytometry and 96-plex immunoassay proteomics on blister biopsy fluid, and spatial transcriptomics and proteomics on FFPE tissues to determine the cellular composition of CLE interface dermatitis. Quantification of cellular infiltrates from blister biopsies revealed significant increases in HLA-DR+ antigen presenting cell populations (p<0.0021 lesional vs nonlesional, p<0.0006 nonlesional vs healthy) as well as CD8+ T cells (p<0.0166 lesional vs healthy, **Fig 2A**). Querying the 96-plex immunoassay proteomics dataset for cell type specific proteins identified increased T cell (CD5, CD8A) and APC markers (CD302) in both lesional and nonlesional blister fluid (**Fig 2B**). These surface markers are likely shed due to the presence of increased matrix metalloproteinases such as ADAM15 (**Fig 2B**). Additional significant protein biomarkers are presented in **Table S4**.

**Fig. 2.**
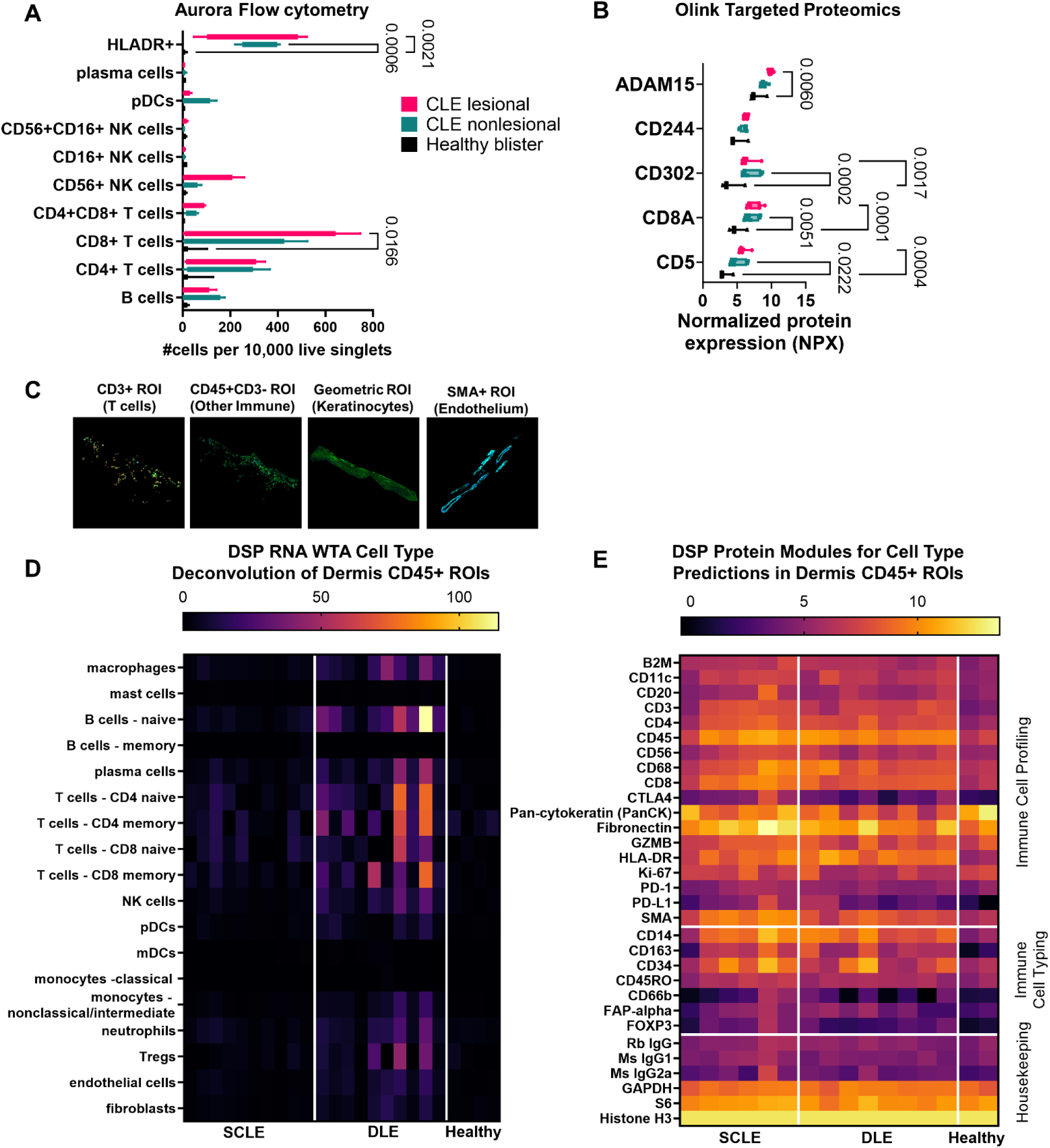
Characterizing immune cell subsets in CLE interface dermatitis reveals significant increases in HLA-DR+ cells and CD8+ T cells. (**A**) Aurora flow cytometry for cell lineage markers in CLE lesional and nonlesional skin compared to healthy blisters. Enumeration of cellular infiltrates were normalized to 10,000 live singlet cells. One-way ANOVA with Tukey’s post hoc tests revealed significant increases in HLA-DR+ cells and CD8+ T cells compared to healthy control blisters, with trends towards increased B cells, CD4+ T cells and NK cells. pDCs were trending higher in nonlesional blister fluid. (n=3 healthy and 4 CLE patients) (**B**) 96-plex immunoassay targeted proteomics for cell lineage markers in blister fluid. T cell markers (CD5, CD8A) and antigen presenting cell markers (CD302) were also significantly elevated in CLE blister fluid compared to healthy controls. NK marker CD244 was not significantly elevated, matching flow cytometry findings. Note that shedding of surface CD molecules is likely due to the presence of proteases such as ADAM15. (n=7 CLE lesional, 7 CLE nonlesional from 4 donors and 3 healthy blisters) (**C**) Examples of Regions of interest (ROIs) chosen for spatial analysis of FFPE CLE biopsies. We used cell masking approaches for CD3+ T cells, CD45+ immune cells, geometric autofluorescent keratinocytes/epidermis and SMA+ endothelium. (**D**) Cell type deconvolution in the CD45+ dermal ROIs from the WTA dataset was calculated using NanoString’s prediction module. We noted increases in T cells and B cells, as well as neutrophils which were not assessed by flow cytometry due to neutrophil death upon extraction from skin. (n=8 DLE, 6 SCLE and 2 healthy margin CD45+ ROIs) (**E**) Spatial proteomics was assessed for the immune cell profiling and immune cell typing modules to predict the presence of immune cells. Housekeeping proteins are depicted in the bottom module. (n=95 ROIs assessed from 4 DLE, 4 SCLE from 3 donors and 3 healthy margin controls for protein DSP)

Next, we examined our cell type regions of interest (ROIs) in the spatial datasets. CD3+CD45+ masks were used to enrich for T cells, CD45+CD3-masks to enrich for other immune cells, geometric autofluorescent ROIs to enrich for keratinocytes and SMA+ ROIs to enrich for endothelium (**Fig 2C**). We were not able to capture many ROIs from endothelium in cross section, therefore we focused our analyses on the T cell, immune cell and keratinocyte ROIs. NanoString GeoMX software deconvolution of the immune cell ROIs in the WTA dataset revealed increases in many immune cell types in the dermis including B cells, T cells, dendritic cells and monocytes/macrophages as assessed by cell-type specific genes (**Fig 2D**). Examination of protein-level expression of CD surface markers using the immune cell profiling and immune cell typing protein modules confirmed CD20+ B cell, CD14+ and CD68+ monocyte/macrophage expression (**Fig 2E**). Taken together, the predominant cell types comprising CLE interface dermatitis are HLADR+ myeloid cells and CD8+ T cells.

### Examining differences in discoid versus subacute cutaneous lupus erythematosus reveals unique gene and immune checkpoint expression

One of the remaining questions in the field of cutaneous lupus is what immunologic differences underlie the pathogenesis of discoid lupus erythematosus (DLE) versus subacute cutaneous lupus erythematosus (SCLE). To this end, we separated out the CLE subtypes and performed both transcriptomic and proteomic analyses comparing the 2 entities, including donor as a correction variable. Spatial transcriptomic analysis of DLE vs. SCLE skin detected 19 DEGs in keratinocyte ROIs using a log2 fold change cutoff of +/- 1 and p value cutoff of 0.05 (**Fig 3A & B**). Pathway analysis of these genes revealed terms related to G protein coupled receptors (GPCRs) enriched in SCLE versus terms involved in RNA processing and interferon signaling in DLE (**Fig 3C**). We also performed hypothesis testing by querying specific subsets of genes across conditions. Examining calpains, which were recently described as a treatment target for photosensitivity^12^, revealed that *CAPN1* was enriched in SCLE, whereas *CAPN3* was enriched in DLE (**Fig 3D**). Analysis of CD45+ ROIs detected 27 DEGs in DLE vs SCLE (**Fig 3E & F**). Pathway analysis of these genes revealed keratinization was enriched in SCLE, whereas interferon alpha/beta signaling and rRNA processing terms were enriched in DLE (**Fig 3G**). Of note, the keratinization term includes immune-related genes such as calpains and cathepsins. Examining IL-1 Receptor Associated Kinases (IRAKs), which were recently described as a treatment target for skin inflammation^13^, revealed *IRAK1* was enriched in DLE, whereas *IRAK4* was enriched in SCLE immune cells (**Fig 3H**). Analysis of CD3+ ROIs detected 4 DEGs in DLE vs SCLE (**Fig 3I & J**). Since so few genes were above a log2 fold change cutoff of 1, we performed analysis of differentially expressed proteins in place of pathway analysis. Interestingly, the immune inhibitory family member B7-H3 was uniquely upregulated in SCLE T cells (**Fig 3K**). Examining Janus kinases (JAKs), which have been used for multiple inflammatory skin disorders, revealed *TYK2* is enriched in both DLE and SCLE T cells compared to healthy (**Fig 3L**). Taken together, these data suggest that nuances in immune cell and keratinocyte signaling may underlie DLE versus SCLE pathogenesis, and that some treatment targets such as B7-H3, calpains or IRAKs may be subtype specific, while others, like TYK2, are shared.

**Fig. 3.**
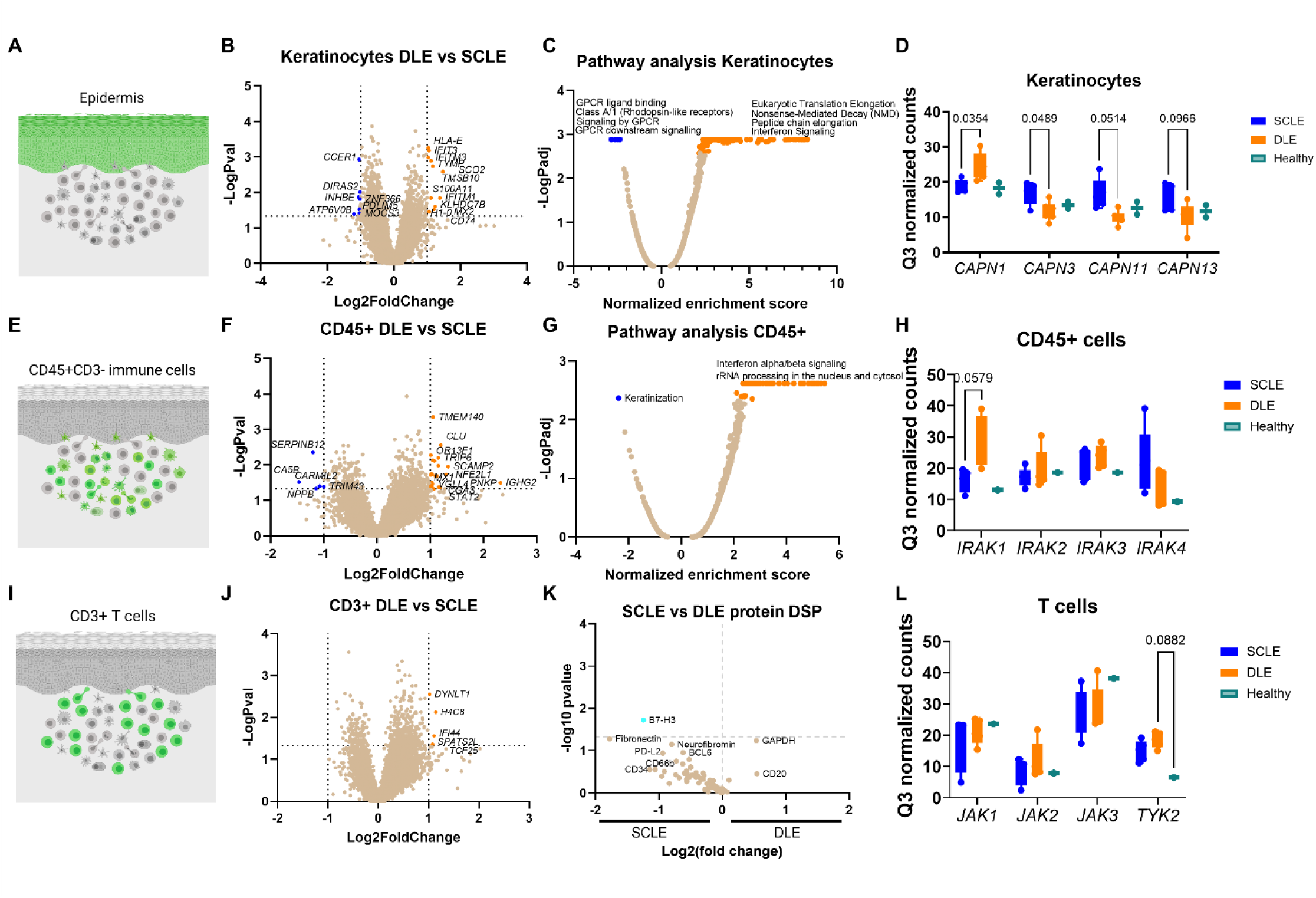
Examining differences in DLE versus SCLE skin biopsies reveals biological differences that may underlie pathogenesis and/or susceptibility to systemic disease. (**A**) Example keratinocyte/epidermal region of interest (ROI) cartoon. (**B**) Volcano plot of the keratinocyte ROI in DLE vs SCLE biopsies. (n=5 SCLE vs 6 DLE keratinocyte ROIs) (**C**) Pathway analysis of the keratinocyte ROI in DLE vs SCLE biopsies as performed in GeoMX software. (**D**) Calpain gene family expression in keratinocytes reveals differences in DLE vs SCLE. (two-way ANOVA with Tukey’s posttests significant or trending, as indicated) (**E**) Example CD45+ ROI cartoon. (**F**) Volcano plot of the CD45+ ROI in DLE vs SCLE biopsies. (n=5 SCLE vs 5 DLE CD45+ ROIs) (**G**) Pathway analysis of the CD45+ ROI in DLE vs SCLE biopsies. (**H**) IL-1 Receptor Associated Kinase (IRAK) gene family expression in CD45+ cells reveals differences in DLE vs SCLE. (two-way ANOVA with Tukey’s posttests significant or trending, as indicated) (**I**) Example CD3+ ROI cartoon. (**J**) Volcano plot of the CD3+ ROI RNA DSP in DLE vs SCLE biopsies. (n=5 SCLE vs 5 DLE CD3+ ROIs) (**K**) Volcano plot of the CD3+ ROI protein DSP in DLE vs SCLE biopsies. (n= 11 DLE, 6 SCLE and 3 healthy CD3+ ROIs) The immune checkpoint B7-H3 is unique to SCLE and may serve as a novel treatment target. (**L**) Janus kinase (JAK) gene family expression in DLE vs SCLE T cells reveals the treatment target *TYK2* is trending higher in both DLE and SCLE. (n=5 SCLE vs 5 DLE CD3+ ROIs, two-way ANOVA with Tukey’s posttests significant or trending, as indicated) Cartoons created with Biorender.com

### Keratinocytes at the inflammatory and lesional sites shape chemotactic signals for leukocyte recruitment

We next asked whether we could identify immune signaling pathways in keratinocytes that could lead to the initiation of inflammation. We selected regions of interest above interface dermatitis that exhibit hyperkeratosis and/or acantholysis in overlying keratinocytes. Comparing keratinocytes that were above interface dermatitis (proximal) versus distal to an infiltrate (**Fig 4A**) revealed upregulation of key chemokines including *Cxcl9* and *Cxcl10* in keratinocytes proximal to the immune infiltrate (**Fig 4B**). 96-plex immunoassay proteomic data in lesional and nonlesional blister fluid was compared to the RNA spatial dataset, and demonstrated that CXCL9 and CXCL11 proteins are significantly upregulated in lesional blister fluid (**Fig 4C**). Additionally, increased levels of chemokine ligands CCL8 and CXCL6 in lesional versus nonlesional blister fluid were identified (**Fig 4C**). We re-queried the WTA dataset to analyze potential innate immune pathways that could lead to chemokine expression and found increased *AIM2* expression (**Fig 4D**). Querying specific members of the AIM2 pathway in the 96-plex immunoassay dataset identified significant increases in caspase 8 ^14^ and the IL1 family member IL18 ^15^ in lesional skin relative to both healthy and nonlesional samples (**Fig 4E & F**). 96-plex immunoassay also identified significant increases in IFNL1 and IFNG protein levels in lesional CLE skin, which are upstream of CXCL9 and CXCL11 (**Fig 4G & H**). Of note, IL18 induces CXCL6 expression in murine lung^16^, and CCL8 is optimally induced by combinations of IL1, IFNG and dsRNA^17^. Taken together, these data suggest that keratinocyte injury, as evidenced by hyperkeratosis and acantholysis visible on histology, can induce AIM2-caspase 8-IL18 pathways and IFN pathways^18^ to turn on chemokines including CCL8, CXCL6, CXCL9 and CXCL11 for recruitment of immune cells in CLE.

**Fig. 4.**
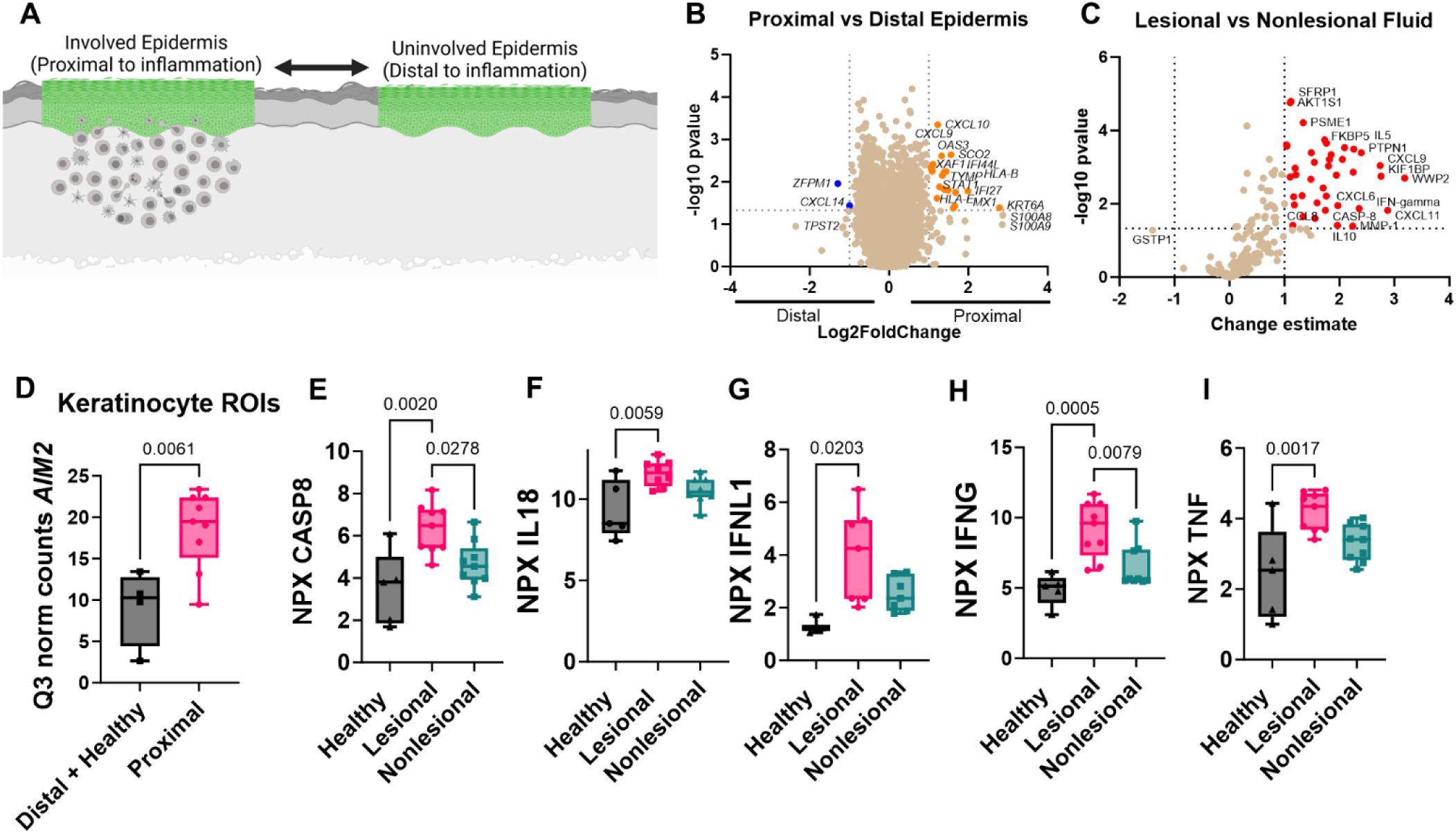
Examining proximal versus distal keratinocytes, and lesional versus nonlesional blister fluid, reveals pathways that may be required for initiation of chemokine cascades for leukocyte recruitment to form interface dermatitis. (**A**) Schematic of proximal versus distal analysis. Created with Biorender.com. (**B**) Volcano plot of DSP WTA data for proximal versus distal epidermis. (n=9 proximal and 4 distal plus healthy margin keratinocyte ROIs) (**C**) Examination of lesional versus nonlesional blister fluid in 96-plex immunoassay as performed with NPX software. (n= 9 lesional and 9 nonlesional blister fluid samples from 6 patients). (**D**) Box plot of *AIM2* expression in keratinocytes from distal plus healthy margins versus proximal keratinocytes from Q3 normalized counts in the WTA dataset. (n=4 healthy plus distal and 9 proximal ROIs) (**E**) Caspase 8, (**F**) IL-18, (**G**) IFNL1 and (**H**) IFNG were elevated in CLE blisters compared to healthy controls in the 96-plex immunoassay dataset (Normalized Protein eXpression, NPX; panels E, F, H & I: n=9 lesional and 9 nonlesional blister fluid samples from 6 patients, and 5 healthy controls run in the 96-plex immunoassay inflammation panel; panel G: n=7 lesional and 7 nonlesional blister fluid samples from 4 patients, and 3 healthy controls run in the 96-plex immunoassay neuroexploratory panel).

### Combining proteomic analysis of blister fluid with spatial transcriptomics of skin tissue reveals cell-specific expression patterns of chemokine ligands in CLE skin

We^19–21^ and others^9,22,23^ have demonstrated upregulation of chemokine ligands in CLE by RNA sequencing and microarray. 96-plex immunoassay proteomics identified significant increases in many chemokine ligands at the protein level, including CXCR3 family, CCL8, and CXCL6 chemokines (**Fig 5A**). Chemokines were fairly stable over time, as determined by repeat blistering of one of our donors approximately 6 months apart (**Fig S5**). Querying the WTA dataset revealed cell-specific context expression of these ligands which differed by both ROI type and disease state (**Fig 5B**). Specifically, *Ccl8* was expressed highly in the CD45+ ROI from DLE biopsies, whereas *Ccl25* was expressed in SCLE biopsies, indicating a different cellular context of these signals. In contrast, *Cxcl6* was expressed by both keratinocytes and immune cells. *Cxcl9/10/11* were expressed mainly by the immune system, with the highest upregulation in DLE, which fits with prior literature describing an IFNG “node” in DLE^9^. Taken together, these data help to map cellular sources of chemokines, which in turn provides insights into the order in which they are produced, whether by skin resident immune cells, keratinocytes or infiltrating immune cells.

**Fig. 5.**
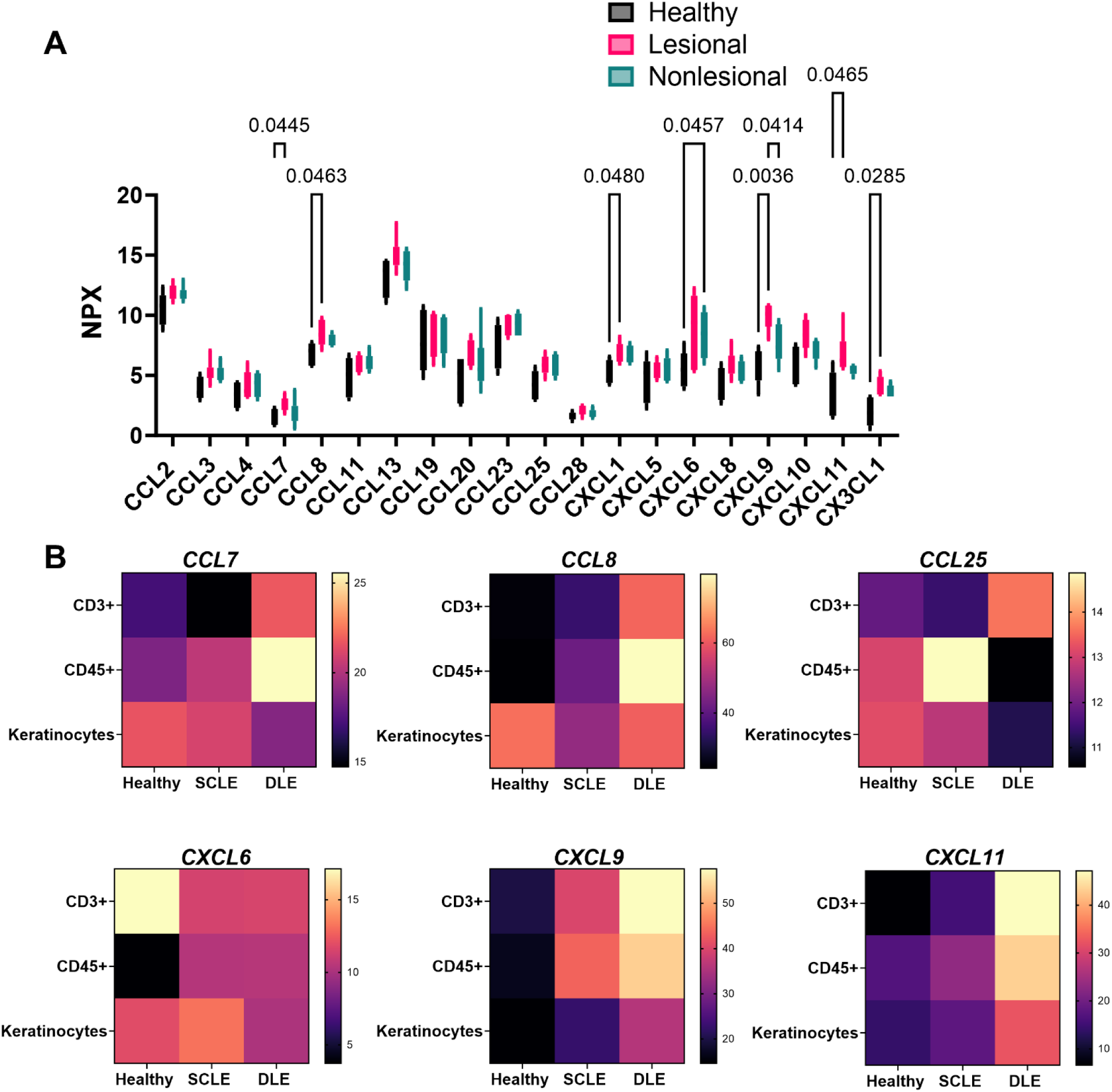
Combining proteomics and spatial transcriptomics using a cell masking approach allows assignment of biomarkers and chemokine ligand-receptor pairs governing leukocyte recruitment to the skin during CLE. (**A**) We queried the 96-plex immunoassay datasets for all available chemokine measurements. Significant differences were observed in the levels of CCL7, CCL8, CXCL1, CXCL6, CXCL9, CXCL11 and CX3CL1 in the 96-plex immunoassay interstitial skin fluid analysis (two-way ANOVA with Tukey’s post tests significant as indicated; n=5 healthy blisters, and 6 matched CLE lesional and nonlesional blisters). **B**) Heatmaps illustrate the cellular sources of chemokines in the WTA DSP analysis for healthy, SCLE, and DLE samples. (n=41 ROIs assessed from 4 DLE, 3 SCLE, and 2 healthy margin control biopsies for WTA DSP)

### Functional chemotaxis assays recapitulate CXCR3-mediated T cell migration and reveal CXCR1-sensitive CD14+CD16+ and CCR2-sensitive CD14+ populations

To determine the functional consequences of our data, blood was drawn from healthy and CLE donors and used in chemotaxis assays, focusing on the ligands we identified in the 96-plex immunoassay dataset (CCL8, CXCL6, CXCL9, CXCL11; **Table S5**). Flow cytometry was also performed on input cells prior to migration to determine distribution of the relevant chemokine receptors (antibody information & chemokine information in **Table S6 & Table S7**). CCL8 binds to CCR1/2/3/5; CXCL6 binds to CXCR1/2, and CXCL9 and CXCL11 both bind to the receptor CXCR3. CCR3 is typically expressed on eosinophils, so we did not assess this receptor, but instead assessed CCR2/5 on monocytes which are the high affinity CCL8 receptors. Cells were gated on live singlets and were identified with major lineage markers (e.g. CD3+ T cells, CD19+ B cells, CD56+ NK cells). Both lupus and healthy donor T cells expressed CXCR3 (**Fig 6A & B**) and were able to migrate well towards CXCL11 (**Fig 6C**). However, lupus donor T cells only exhibited a 2-fold increase in migration over media baseline in lupus patient samples, compared to an average 5-fold increase in migration over media baseline in healthy samples (**Fig 6D**). We hypothesize this may be due to maintenance therapy in lupus donors. Notably, CXCL9 and CXCL11 are off in healthy skin, and turned on in lupus skin, exemplifying how T cells are poised to respond to IFN-inducible chemokines.

**Fig. 6.**
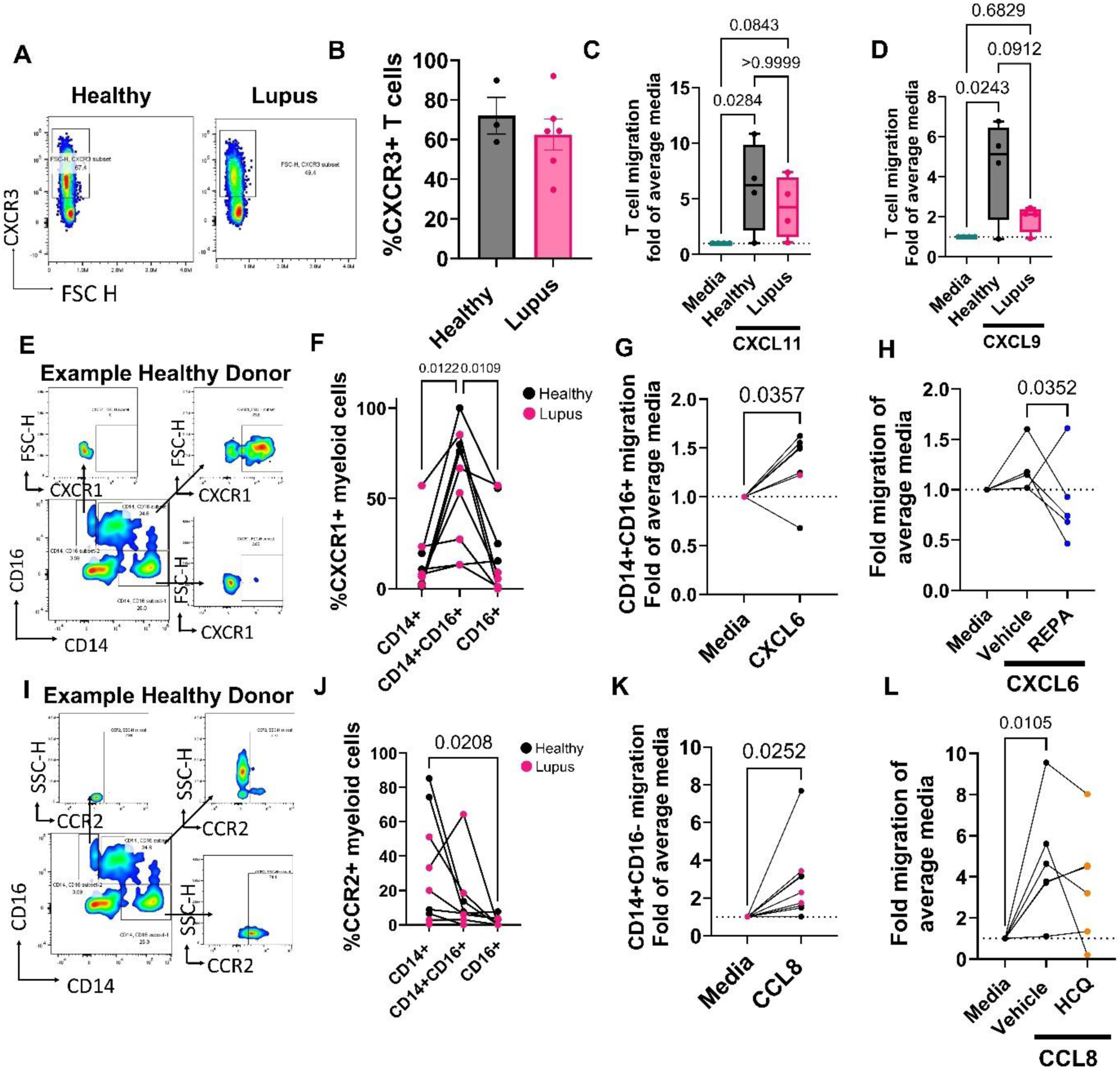
Functional chemotaxis assays reveal that T cells are poised to respond to CXCR3 ligands, whereas CD14+CD16+ myeloid cells are poised to respond to CXCL6 via CXCR1 and CD14+CD16-myeloid cells are poised to respond to CCL8 via CCR2. (**A**) Example flow staining and (**B**) quantification of CXCR3 on CD3+ T cells (n=3 healthy and n=6 CLE, t test ns). (**C**) Chemotaxis assay reveals both healthy and lupus T cells migrate towards CXCL11 (n=4 healthy and 4 lupus donors; one-way ANOVA with significant posttests as indicated). (**D**) Chemotaxis assay reveals higher migration in healthy donor T cells than lupus donor T cells towards CXCL9. (n=4 healthy and 4 lupus donors; one-way ANOVA with significant posttests as indicated). (**E**) Example flow staining of myeloid cells demonstrating CXCR1 expression in CD14+, CD14+CD16+ and CD16+ gates. (**F**) Quantification of CXCR1+ cells demonstrates CD14+CD16+ cells express more CXCR1 than their counterparts. (n=4 healthy and 5 lupus donors; p=0.0122 and 0.0109 by repeated measures one way ANOVA matched by donor). (**G**) Chemotaxis assay reveals directed migration of CD14+CD16+ myeloid cells in all but one healthy donor (n=6 healthy and 1 lupus donor; paired t test p=0.0357). (**H**) Chemotaxis assay reveals pretreatment of PBMCs from healthy donors with the CXCR1 inhibitor reparixin (REPA) inhibits migration towards CXCL6. (n=5 healthy donors; one way ANOVA with significant posttests as indicated). (**I**) Example flow staining of myeloid cells demonstrating CCR2 expression in CD14+, CD14+CD16+ and CD16+ gates. (**J**) Quantification of CCR2+ myeloid cells demonstrates CD14+CD16-cells are trending towards more CCR2 expression than their counterparts (n=4 healthy and 5 lupus donors; p=0.0208 by one way ANOVA with significant posttests as indicated). (**K**) Chemotaxis assay demonstrated that CD14+CD16-cells migrate towards CCL8. (n=6 healthy and n=3 lupus donors; paired t test p=0.0252). (**H**) Chemotaxis assay reveals pretreatment of PBMCs from healthy donors with hydroxychloroquine (HCQ) inhibits migration towards CCL8 (n=6 healthy donors; one way ANOVA with significant posttests as indicated).

We next assessed CXCL6 and its cognate receptor CXCR1^24^, which has been reported to be expressed on skin NK cells^25^ and is important for their recruitment into tumors^26^. NK cells from both healthy and lupus donors expressed CXCR1 (**Fig S6A**), though there was donor variability for migratory capacity towards the CXCL6 ligand. Like T cells, NK cells migrated to CXCL9 and CXCL11 (**Fig S6B & C**). We also assessed B cell migration, which we observed to be dependent on CXCR5-CXCL13 (**Fig S6D-F**). These findings fit with current literature describing NK cell and B cell migration pathways.

Last, we assessed chemokine expression on myeloid cells, which likely constitute the HLA-DR+ population we noted to be increased in blister biopsies. CD14+CD16+ myeloid cells were recently identified in nonlesional CLE skin by Billi et al using 10X spatial transcriptomics and were hypothesized to be the first responders to skin injury/inflammation in lupus^27^. We gated live, single, CD45+CD3-CD19-CD56-cells (i.e. non-T/B/NK) followed by CD14 and CD16 expression (**Fig S7**) to confirm these cells are present in CLE blood and skin by flow cytometry (**Fig S8**). We examined chemokine receptor expression in CD14+CD16-, CD14+CD16+ and CD14-CD16+ myeloid cells and noted significant increases in CXCR1 expression on CD14+CD16+ myeloid cells (**Fig 6E &F**). Total number of monocytes were higher at baseline in healthy versus lupus patients, which we hypothesize may be due to maintenance therapy in lupus patients. All CD14+ myeloid cells, but particularly CD14+CD16+ myeloid cells, migrated towards CXCL6, providing a possible chemokine receptor:ligand pair that mediates their entry into the skin (**Fig 6G**). To assess a potential inhibitor of this pathway, we treated healthy donor PBMCs with the CXCR1/2 inhibitor reparixin (REPA) and assessed migration towards CXCL6. REPA treatment reduced migration in all but one donor, who also paradoxically did not migrate well towards CXCL6 (**Fig 6H**). Taken together, these data identify a potential inhibitor of CD14+CD16+ cell entry into lupus skin, which could prevent new lesion formation.

In contrast to CXCR1, the CCL8 receptor CCR2 was enriched on CD14+CD16-myeloid cells (**Fig 6I & J**), but not the CCR5 receptor (**Fig S9**). CD14+CD16-myeloid cells migrated towards CCL8 (**Fig 6K**). To understand the potential effects of hydroxychloroquine (HCQ) on this pathway, we pretreated healthy PBMCs and subjected them to chemotaxis assays. HCQ treatment reduced CCL8 directed migration by CD14+CD16-monocytes (**Fig 6L**).

Taken together, these data exemplify how disease states can influence chemokine ligand levels and influence timing and recruitment of specific immune cell populations to the tissue, namely through keratinocytes following injury and subsequently through recruited immune cell populations. We propose the model that keratinocyte injury induces activation of AIM2-related pathways that turn on CXCL6 and CCL8 expression to recruit myeloid cells, which in turn potentiate IFN-related signaling pathways to recruit T cells and other lymphocytes via CXCR3 ligands (**Graphical abstract**).

## DISCUSSION

In this study, we sought to analyze chemokine orchestrators of T cell rich interface dermatitis in CLE. Chemokines can be released by keratinocytes in response to UV light, infection, or other environmental triggers^28^. These chemokines can then recruit immune cells to the skin, which can lead to inflammation. One of the most highly upregulated chemokine families in CLE lesions is the CXCR3 chemokine family^6,29,30^. CXCR3 binds to its interferon (IFN)-inducible ligands CXCL9, CXCL10 and CXCL11 to mediate leukocyte migration and function^31^. IFN signaling has been postulated to drive the pathogenesis of all subtypes CLE^32^. Further, CLE is a photosensitive disorder, and UV light induces upregulation of IFN to amplify CXCR3 ligand production in keratinocytes^18,33,34^, linking the environmental insult to the recruitment of pathogenic immune cells in the context of CLE. The interaction between CXCR3-expressing T-cells and its ligands CXCL9, CXCL10 and CXCL11 have been associated with tissue damage in several CLE subtypes^8^. Examining chemokine ligand expression in our 96-plex immunoassay dataset and in our ROIs, as well as chemokine receptor expression in our flow cytometry data, confirmed expression of the CXCR3 family and also identified potential novel mediators of recruitment in CLE. Based on 96-plex immunoassay and flow cytometry data, we hypothesize that CXCR1 is a key receptor for CXCL6 on proinflammatory CD14+CD16+ APCs, and could be a viable treatment target for CLE, especially given the role of CXCL6 in fibrosis which occurs in DLE and other forms of chronic CLE^35^ as well as in renal interstitial fibrosis^36^. CXCL6 is a particularly interesting target immunologically, given that it may be upregulated by double stranded RNA or IL1𝝱^37^. This is important, as we know from work by Rodriguez-Pla et al. that IFN priming alone is not sufficient to activate migratory programs in lupus monocytes^38^, and from work by Billi et al. that CD14+CD16+ monocyte/dendritic/myeloid cell populations are among the first cells recruited to the skin in CLE^27^. Further studies are needed to understand epigenetic factors that result in increased CXCR1 expression in CD14+CD16+ myeloid cells.

The 96-plex immunoassay dataset provides insight into novel protein biomarkers of CLE beyond chemokine ligands that have interesting implications in immunopathogenesis, including Flt3L, HGF, CD40 and CEACAM3. Flt3L is a dendritic cell growth and survival factor, and has been previously described in SLE^39^. A recent study demonstrated generation of tolerogenic conventional DCs in SLE patients undergoing mesenchymal stem cell therapy^40^. Thus, it is possible that the upregulation of Flt3L in skin is a compensatory mechanism. HGF, which as noted above can increase CXCR3 expression on T cells^11^, is higher in SLE patients than healthy controls^41^ and in patients with higher disease activity^42^. HGF administration can prevent lupus nephritis in mice^43^, potentially indicating a compensatory mechanism for HGF in CLE. CD40 overexpression causes autoimmune skin disease in mice^44^, and increased CD40 expression has been reported in IHC of CLE lesions^45^. Genome wide differential expression analysis identified CEACAM3 as a potential gene involved in the pathogenesis of lupus and lupus nephritis^46^, though its role in skin disease has yet to be elucidated.

We also found differences in DLE and SCLE that have not previously been appreciated by using spatial technologies. For example, DLE lesional skin T cells express less B7-H3 than SCLE lesional skin as assessed by protein DSP. B7-H3, also known as CD276, is an immune checkpoint within the B7 molecular family that fine tunes immune responses^47^ and is currently being explored as a cancer immunotherapy target ^48^. B7-H3 plays a pivotal role in mediating the suppressive effects of DCs on T cells and downregulating Th1 responses^49^. Therefore, a relative reduction in B7-H3 in DLE as compared to SCLE fits with previously described IFNG signaling nodes identified in DLE by bulk RNA sequencing^9^. Further studies are needed to assess the DEGs for DLE vs SCLE ROI comparisons that were identified in the WTA dataset.

In addition to insights into immune cell recruitment and potential novel biomarkers, our datasets identify potential treatment targets. Reparixin and the related drug ladarixin are CXCR1 inhibitors that are currently in clinical trials for cancer^50^. It would be interesting to test a topical formulation for CLE for prevention of new or spreading lesions. A calpain 1 inhibitor was recently demonstrated to prevent photosensitivity in animal models of UVB exposure ^12^. We noted elevated *CAPN1* in SCLE keratinocytes, which makes this another attractive topical treatment to test. TYK2 inhibitors are currently in clinical trials for SLE^51^, and several case reports have noted improvement in CLE^52^. We found elevated *TYK2* in T cell ROIs from both DLE and SCLE patients. A small molecule inhibitor of IRAK4 was recently described as a treatment target for psoriatic skin inflammation ^13^. We identified elevated *IRAK4* in DLE, and *IRAK1* in SCLE immune cells. Given that both TYK2 and IRAKs were elevated in the immune compartment ROIs, these treatments may require oral rather than topical administration for highest efficacy. We hope that these data inspire future clinical trials repurposing drugs for CLE, as well as studies testing topical versus oral routes of administration for more advanced compounds.

While our study provides valuable insights into the immunopathogenesis of CLE, it is essential to acknowledge the study’s limitations. First, our research predominantly focuses on a specific subset of chemokine ligand-receptor pairs and immune cell interactions that we were able to measure with current methodologies. CXCR1 is a classical receptor for neutrophil recruitment to inflamed tissues, and ligation of CXCR1 can enhance NET formation^53,54^. We were not able to recover neutrophils from blister fluid, and opted to use Ficoll gradients, which remove the bulk of granulocytes, to isolate PBMCs for chemotaxis assays. Future investigations should include methodologies to specifically assess neutrophils and other granulocytes. Moreover, a longitudinal approach with serial assessments would offer a more subtle understanding of disease progression over time. Finally, our findings in different CLE subtypes and patient populations should be examined further with larger patient cohorts. Future studies exploring longitudinal analyses with more patients, and comparing myeloid cells in the skin of patients with response or resistance to antimalarials, are warranted to confirm and expand upon these findings. In-depth mechanistic studies and the development of targeted interventions based on the identified chemokine and other pathways holds promise for improving therapeutic strategies in CLE.

## List of Supplementary Materials

Supplemental Materials & Methods

Fig S1. Quality Control for Digital Spatial Profiling (DSP) Whole Transcriptome Atlas (WTA) spatial transcriptomics.

Fig S2. Quality Control and Pathway Analysis for Regions of Interest (ROIs).

Fig S3. Validation of DSP dataset using historical microarray dataset and a cancer transcriptome atlas (CTA) dataset.

Fig S4. Validation of DLE dataset ROIs as compared to CTA.

Fig S5. Stability of chemokines in lesional and nonlesional samples over time.

Fig S6. Assessment of NK cell and B cell migration.

Fig S7. Flow gating strategy for assessing myeloid populations.

Fig S8. Examination of CD14+CD16+ cells in blister biopsies

Fig S9. CCR5 is not enriched on CD14 vs CD16 expressing myeloid cells.

Table S1. Blister biopsy and blood donation patient info & characterization (UMass Chan).

Table S2. Archival CLE biopsies used in WTA Digital Spatial Profiling (UMass Chan).

Table S3. Validation archival CLE biopsies used in CTA Digital Spatial Profiling (Yale).

Table S4. 96-plex immunoassay DEPs calculated by NPX software and 2-way ANOVA.

Table S5. Chemotaxis and chemokine receptor staining blood donor information (UMass Chan and Dartmouth Hitchcock).

Table S6. Antibody information & RRIDs.

Table S7. Chemokine information.

## Acknowledgments

We thank those who donated tissue for this study. We thank Celia Hartigan for clinical research support, and Jonathan Reichner for feedback on our manuscript. We thank Alaina Ryan and the Comparative Pathobiology and Genomics Shared Resources (“CPGSR”) in Cummings School of Veterinary Medicine at Tufts University for running the protein DSP plate in the MAX machine. We used Biorender.com for illustrations. SS used ChatGPT to correct English grammar in the paragraphs he wrote for the first draft of the manuscript, and JMR edited for clarity and brevity as a native English speaker.

## Contributor statement

Conceptualization: JMR

Methodology: JMR, MR, SS, YL, CEB, JEH

Investigation: MAR, EK, JGDeL, JPS, AA, KA, LZ, ML, KD, EMacD, MDV

Validation: SS, UYA, MDV, SSher, MD, MR

Formal analysis: SS, NSH, KA, JEL, HSR, UYA, RL, JMR Resources: AD, MR, CEB, JBW, SSG, JEH, MDV, JMR

Data Curation: SS, NSH, KA, JMR, JV, PR

Visualization: SS, NSH, KA, JEL, UYA, MD, JMR Supervision: JMR, MR

Project administration: JMR Funding acquisition: JMR

Writing - Original Draft: SS, NSH & JMR

Writing - Review & Editing: all authors

## Data availability statement

Datasets related to this article have been deposited in National Center for Biotechnology Information’s Gene Expression Omnibus (GEO) *(72)* and are accessible through GEO Series accession numbers GSE182302 for 96-plex immunoassay data, and GSE182825 for spatial transcriptomics. Flow data has been deposited on FlowRepository under accessions FR-FCM-Z4PL, FR-FCM-Z4PM, FR-FCM-Z4PN, FR-FCM-Z4PQ, FR-FCM-Z4PX, FR-FCM-Z6UN, FR-FCM-Z7ZP, FR-FCM-Z7ZQ for blister biopsies and blood immunophenotyping.

## Ethics statement

This study was performed with internal review board (IRB) approved protocols at the UMass Chan Medical School (H-14848 and H00021295 for fresh blood and skin tissue; H00020503 for archival skin tissue), Yale University Institutional Review Board (Human Investigative Committee no. 15010105235 for archival skin tissue), and/or Dartmouth Hitchcock Medical Center (STUDY02001542 for PBMCs), and all samples were obtained with written informed consent and were de-identified before use in experiments.

## Bulleted statements

- *What is already known about this topic?* Cutaneous lupus erythematosus (CLE) is a rare, chronic autoimmune connective tissue disorder. Nonlesional skin was recently reported to be enriched in CD14+CD16+ inflammatory myeloid cells.
- *What does this study add?* We mapped chemokine orchestrators of interface dermatitis in lupus using spatial approaches on archival tissue and confirmed with fresh tissues, to understand how keratinocytes, myeloid cells, T cells and other leukocytes form the interface dermatitis reaction.
- *What is the translational message*? The translational message of our findings is that, even on maintenance therapy, CLE patient skin is enriched for both T cells and antigen presenting cells. Keratinocyte injury activates innate immune pathways that ultimately induce chemokines. These chemokines recruit leukocytes, which begets additional cascades of chemokines. Specifically, we show that the CD14+CD16+ inflammatory myeloid cells respond to CXCL6, which is upregulated at the protein level in CLE lesional interstitial skin fluid.
- *What are the clinical implications of this work*? Clinically, our findings have implications for drug repurposing and future clinical trials. Namely, the CXCL6/CXCR1 inhibitor reparixin and its derivatives, which are currently used in oncological applications, could be tested in topical formulations for CLE.

## References

1 Garelli CJ, Refat MA, Nanaware PP, et al. Current Insights in Cutaneous Lupus Erythematosus Immunopathogenesis. Front Immunol 2020; 11:1353.

2 Schultz HY, Dutz JP, Furukawa F, et al. From pathogenesis, epidemiology, and genetics to definitions, diagnosis, and treatments of cutaneous lupus erythematosus and dermatomyositis: a report from the 3rd International Conference on Cutaneous Lupus Erythematosus (ICCLE) 2013. J Invest Dermatol 2015; 135:7–12.

3 Reich A, Werth VP, Furukawa F, et al. Treatment of cutaneous lupus erythematosus: current practice variations. Lupus 2016; 25:964–72.

4 Chang AY, Werth VP. Treatment of cutaneous lupus. Curr Rheumatol Rep 2011; 13:300–7.

5 Stannard JN, Kahlenberg JM. Cutaneous lupus erythematosus: updates on pathogenesis and associations with systemic lupus. Curr Opin Rheumatol 2016; 28:453–9.

6 Wenzel J, Zahn S, Mikus S, et al. The expression pattern of interferon-inducible proteins reflects the characteristic histological distribution of infiltrating immune cells in different cutaneous lupus erythematosus subsets. Br J Dermatol 2007; 157:752–7.

7 Wenzel J, Henze S, Wörenkämper E, et al. Role of the Chemokine Receptor CCR4 and its Ligand Thymus- and Activation-Regulated Chemokine/CCL17 for Lymphocyte Recruitment in Cutaneous Lupus Erythematosus. J Invest Dermatol 2005; 124:1241–8.

8 Méndez-Flores S, Hernández-Molina G, Azamar-Llamas D, et al. Inflammatory chemokine profiles and their correlations with effector CD4 T cell and regulatory cell subpopulations in cutaneous lupus erythematosus. Cytokine 2019; 119:95–112.

9 Berthier CC, Tsoi LC, Reed TJ, et al. Molecular Profiling of Cutaneous Lupus Lesions Identifies Subgroups Distinct from Clinical Phenotypes. J Clin Med Res 2019; 8. doi:10.3390/jcm8081244.

10 Strassner JP, Rashighi M, Ahmed Refat M, et al. Suction blistering the lesional skin of vitiligo patients reveals useful biomarkers of disease activity. J Am Acad Dermatol 2017; 76:847–55.e5.

11 Komarowska I, Coe D, Wang G, et al. Hepatocyte Growth Factor Receptor c-Met Instructs T Cell Cardiotropism and Promotes T Cell Migration to the Heart via Autocrine Chemokine Release. Immunity 2015; 42:1087–99.

12 Doleckova I, Vidovic T, Jandova L, et al. Calpain inhibition protects against UVB-induced degradation of dermal-epidermal junction-associated proteins. J Invest Dermatol 2024. doi:10.1016/j.jid.2024.02.020.

13 Lavazais S, Jargosch M, Dupont S, et al. IRAK4 inhibition dampens pathogenic processes driving inflammatory skin diseases. Sci Transl Med 2023; 15:eabj3289.

14 Pierini R, Juruj C, Perret M, et al. AIM2/ASC triggers caspase-8-dependent apoptosis in Francisella-infected caspase-1-deficient macrophages. Cell Death Differ 2012; 19:1709–21.

15 Ratsimandresy RA, Indramohan M, Dorfleutner A, Stehlik C. The AIM2 inflammasome is a central regulator of intestinal homeostasis through the IL-18/IL-22/STAT3 pathway. Cell Mol Immunol 2016; 14:127–42.

16 Ishikawa Y, Yoshimoto T, Nakanishi K. Contribution of IL-18-induced innate T cell activation to airway inflammation with mucus hypersecretion and airway hyperresponsiveness. Int Immunol 2006; 18:847–55.

17 Struyf S, Proost P, Vandercappellen J, et al. Synergistic up-regulation of MCP-2/CCL8 activity is counteracted by chemokine cleavage, limiting its inflammatory and anti-tumoral effects. Eur J Immunol 2009; 39:843–57.

18 Skopelja-Gardner S, An J, Tai J, et al. The early local and systemic Type I interferon responses to ultraviolet B light exposure are cGAS dependent. Sci Rep 2020.URL https://www.nature.com/articles/s41598-020-64865-w.

19 Ko W-CC, Li L, Young TR, et al. Gene expression profiling in skin reveals strong similarities between subacute and chronic cutaneous lupus that are distinct from lupus nephritis. J Invest Dermatol 2021. doi:10.1016/j.jid.2021.04.030.

20 Mande P, Zirak B, Ko W-C, et al. Fas ligand promotes an inducible TLR-dependent model of cutaneous lupus-like inflammation. J Clin Invest 2018; 128:2966–78.

21 Garelli CJ, Wong NB, Piedra-Mora C, et al. Shared inflammatory and skin-specific gene signatures reveal common drivers of discoid lupus erythematosus in canines, humans and mice. Current Research in Immunology 2021. doi:10.1016/j.crimmu.2021.03.003.

22 Dey-Rao R, Smith JR, Chow S, Sinha AA. Differential gene expression analysis in CCLE lesions provides new insights regarding the genetics basis of skin vs. systemic disease. Genomics 2014; 104:144–55.

23 Jabbari A, Suárez-Fariñas M, Fuentes-Duculan J, et al. Dominant Th1 and Minimal Th17 Skewing in Discoid Lupus Revealed by Transcriptomic Comparison with Psoriasis. J Invest Dermatol 2014; 134:87–95.

24 Wuyts A, Proost P, Lenaerts JP, et al. Differential usage of the CXC chemokine receptors 1 and 2 by interleukin-8, granulocyte chemotactic protein-2 and epithelial-cell-derived neutrophil attractant-78. Eur J Biochem 1998; 255:67–73.

25 Ottaviani C, Nasorri F, Bedini C, et al. CD56brightCD16(-) NK cells accumulate in psoriatic skin in response to CXCL10 and CCL5 and exacerbate skin inflammation. Eur J Immunol 2006; 36:118–28.

26 Ng YY, Tay JCK, Wang S. CXCR1 Expression to Improve Anti-Cancer Efficacy of Intravenously Injected CAR-NK Cells in Mice with Peritoneal Xenografts. Mol Ther Oncolytics 2020; 16:75–85.

27 Billi AC, Ma F, Plazyo O, et al. Nonlesional lupus skin contributes to inflammatory education of myeloid cells and primes for cutaneous inflammation. Sci Transl Med 2022; 14:eabn2263.

28 Jiang Y, Tsoi LC, Billi AC, et al. Cytokinocytes: the diverse contribution of keratinocytes to immune responses in skin. JCI Insight 2020; 5. doi:10.1172/jci.insight.142067.

29 Flier J, Boorsma DM, van Beek PJ, et al. Differential expression of CXCR3 targeting chemokines CXCL10, CXCL9, and CXCL11 in different types of skin inflammation. J Pathol 2001; 194:398–405.

30 Wenzel J, Wörenkämper E, Freutel S, et al. Enhanced type I interferon signalling promotes Th1-biased inflammation in cutaneous lupus erythematosus. The Journal of Pathology: A Journal of the Pathological Society of Great Britain and Ireland 2005; 205:435–42.

31 Groom JR, Luster AD. CXCR3 ligands: redundant, collaborative and antagonistic functions. Immunol Cell Biol 2011; 89:207–15.

32 Robinson ES, Werth VP. The role of cytokines in the pathogenesis of cutaneous lupus erythematosus. Cytokine 2015; 73:326–34.

33 Meller S, Winterberg F, Gilliet M, et al. Ultraviolet radiation--induced injury, chemokines, and leukocyte recruitment: an amplification cycle triggering cutaneous lupus erythematosus. Arthritis & Rheumatism 2005; 52:1504–16.

34 Scholtissek B, Zahn S, Maier J, et al. Immunostimulatory Endogenous Nucleic Acids Drive the Lesional Inflammation in Cutaneous Lupus Erythematosus. J Invest Dermatol 2017; 137:1484–92.

35 Dai C-L, Yang H-X, Liu Q-P, et al. CXCL6: A potential therapeutic target for inflammation and cancer. Clin Exp Med 2023. doi:10.1007/s10238-023-01152-8.

36 Sun M-Y, Wang S-J, Li X-Q, et al. CXCL6 Promotes Renal Interstitial Fibrosis in Diabetic Nephropathy by Activating JAK/STAT3 Signaling Pathway. Front Pharmacol 2019; 10:224.

37 Wuyts A, Struyf S, Gijsbers K, et al. The CXC Chemokine GCP-2/CXCL6 Is Predominantly Induced in Mesenchymal Cells by Interleukin-1β and Is Down-Regulated by Interferon-γ: Comparison with Interleukin-8/CXCL8. Lab Invest 2003; 83:23–34.

38 Rodriguez-Pla A, Patel P, Maecker HT, et al. IFN priming is necessary but not sufficient to turn on a migratory dendritic cell program in lupus monocytes. J Immunol 2014; 192:5586– 98.

39 Gill MA, Blanco P, Arce E, et al. Blood dendritic cells and DC-poietins in systemic lupus erythematosus. Hum Immunol 2002; 63:1172–80.

40 Yuan X, Qin X, Wang D, et al. Mesenchymal stem cell therapy induces FLT3L and CD1c+ dendritic cells in systemic lupus erythematosus patients. Nat Commun 2019; 10:2498.

41 Robak E, Woźniacka A, Sysa-Jedrzejowska A, et al. Serum levels of angiogenic cytokines in systemic lupus erythematosus and their correlation with disease activity. Eur Cytokine Netw 2001; 12:445–52.

42 Liu Y, Zheng M, Yin W-H, Zhang B. Relationship of serum levels of HGF and MMP-9 with disease activity of patients with systemic lupus erythematosus. Zhejiang Da Xue Xue Bao Yi Xue Ban 2004; 33:340–3, 348.

43 Kuroiwa T, Iwasaki T, Imado T, et al. Hepatocyte growth factor prevents lupus nephritis in a murine lupus model of chronic graft-versus-host disease. Arthritis Res Ther 2006; 8:R123.

44 Mehling A, Loser K, Varga G, et al. Overexpression of CD40 ligand in murine epidermis results in chronic skin inflammation and systemic autoimmunity. J Exp Med 2001; 194:615– 28.

45 Caproni M, Torchia D, Antiga E, et al. The CD40/CD40 ligand system in the skin of patients with subacute cutaneous lupus erythematosus. J Rheumatol 2007; 34:2412–6.

46 AlFadhli S, Ghanem AAM, Nizam R. Genome-wide differential expression reveals candidate genes involved in the pathogenesis of lupus and lupus nephritis. Int J Rheum Dis 2016; 19:55–64.

47 Yi KH, Chen L. Fine tuning the immune response through B7-H3 and B7-H4. Immunol Rev 2009; 229:145–51.

48 Castellanos JR, Purvis IJ, Labak CM, et al. B7-H3 role in the immune landscape of cancer. Am J Clin Exp Immunol 2017; 6:66–75.

49 Suh W-K, Gajewska BU, Okada H, et al. The B7 family member B7-H3 preferentially down-regulates T helper type 1-mediated immune responses. Nat Immunol 2003; 4:899–906.

50 Piro G, Carbone C, Agostini A, et al. CXCR1/2 dual-inhibitor ladarixin reduces tumour burden and promotes immunotherapy response in pancreatic cancer. Br J Cancer 2022; 128:331–41.

51 Morand E, Pike M, Merrill JT, et al. Deucravacitinib, a tyrosine kinase 2 inhibitor, in systemic lupus erythematosus: A phase II, randomized, double-blind, placebo-controlled trial. Arthritis Rheumatol 2023; 75:242–52.

52 Bouché N, Al-Saedy MA, Song EJ. Successful treatment of refractory subacute cutaneous lupus erythematosus with deucravacitinib. JAAD Case Rep 2023; 39:93–5.

53 Teijeira Á, Garasa S, Gato M, et al. CXCR1 and CXCR2 Chemokine Receptor Agonists Produced by Tumors Induce Neutrophil Extracellular Traps that Interfere with Immune Cytotoxicity. Immunity 2020; 52:856–71.e8.

54 Florey OJ, Johns M, Esho OO, et al. Antiendothelial cell antibodies mediate enhanced leukocyte adhesion to cytokine-activated endothelial cells through a novel mechanism requiring cooperation between FcγRIIa and CXCR1/2. Blood 2007; 109:3881–9.

55 Edgar R, Domrachev M, Lash AE. Gene Expression Omnibus: NCBI gene expression and hybridization array data repository. Nucleic Acids Res 2002; 30:207–10.

