## Supplementary material for "Spatial characterization of interface dermatitis in cutaneous lupus reveals novel chemokine ligand-receptor pairs that drive disease": Fig S

**List of Supplementary Materials:**

Supplemental Materials & Methods

Fig S1. Quality Control for Digital Spatial Profiling (DSP) Whole Transcriptome Atlas (WTA) spatial transcriptomics.

Fig S2. Quality Control and Pathway Analysis for Regions of Interest (ROIs).

Fig S3. Validation of DSP dataset using historical microarray dataset and a cancer transcriptome atlas (CTA) dataset.

Fig S4. Validation of DLE dataset ROIs as compared to CTA.

Fig S5. Stability of chemokines in lesional and nonlesional samples over time.

Fig S6. Assessment of NK cell and B cell migration.

Fig S7. Flow gating strategy for assessing myeloid populations.

Fig S8. Examination of CD14+CD16+ cells in blister biopsies

Fig S9. CCR5 is not enriched on CD14 vs CD16 expressing myeloid cells.

Table S1. Blister biopsy and blood donation patient info & characterization (UMass Chan).

Table S2. Archival CLE biopsies used in WTA Digital Spatial Profiling (UMass Chan).

Table S3. Validation archival CLE biopsies used in CTA Digital Spatial Profiling (Yale).

Table S4. 96-plex immunoassay DEPs calculated by NPX software and 2-way ANOVA.

Table S5. Chemotaxis and chemokine receptor staining blood donor information (UMass Chan and Dartmouth Hitchcock).

Table S6. Antibody information & RRIDs.

Table S7. Chemokine information.

**Supplemental Materials & Methods**

**Study design -** The main objective of this study was to characterize chemokine ligand:receptor pairs in CLE to better understand how they orchestrate the interface dermatitis reaction. We set out to answer the question: how are autoreactive T cells recruited to the dermal:epidermal junction in CLE? We hypothesized that, in addition to previously published literature about the CXCR3 chemokine axis in CLE (see for example *(6, 8, 28, 29, 32)*), additional non-redundant chemokine axes were present in the skin to govern immune cell recruitment. As our study progressed, additional literature was published regarding CD14+CD16+ cells in nonlesional CLE skin *(26)*, which we queried in our datasets and performed functional chemotaxis assays. In using -omics approaches, we also sought to provide unbiased analyses that might yield new insights and targets, to perform dataset concordance analyses, to provide protein-level expression data in addition to previously published RNA-level expression data in the CLE field (see for example *(9, 18, 21)*), and to answer sub-questions including: can we identify DEGs and/or DEPs in the different major clinical subtypes of CLE?

**Blister biopsies & blood collection -** Suction blisters (1 cm in diameter) were induced on the skin by using the Negative Pressure Instrument Model NP-4 (Electronic Diversities, Finksburg, MD) as previously described*(10)*. Briefly, the suction chambers were applied to the skin with 10-15 mm Hg of negative pressure and a constant temperature of 40C. After blister formation (30-60min), the blister fluid was aspirated using a 1 mL insulin syringe. Cells within the blister fluid were pelleted at 330 x g for 5-10 minutes for cell staining and the supernatant was collected and frozen for future analysis. The cell pellet was resuspended in FACS buffer (1% FBS in PBS; Sigma Aldrich) and transferred to a FACS tube for flow staining. Venipuncture was performed using heparinized tubes, and peripheral blood mononuclear cells (PBMCs) were isolated following a Ficoll density gradient. Cells were washed once in RPMI (Sigma Millipore) and were resuspended in Cell staining buffer (Biolegend) prior to staining and fixation with Fluorofix buffer (Biolegend). These patient samples are noted in **Table S1** and in the appropriate results sections, and data are deposited on FlowRepository.

**96-plex immunoassay proteomics**: Blister fluid or plasma isolated from heparinized blood was centrifuged at 330 x g for 5 min and aliquoted into 96 well qPCR plates. Samples were shipped to Olink Proteomics Inc. (Cambridge, MA) for analysis in the inflammation and neuroexploratory panels. An additional 2 patients and 2 controls were run in the inflammation panel at the UMass Chan Integrative Biomarker Core. Blister fluids were analyzed using Olink Inflammation and Neuro Exploratory panels in a high throughput proteomics platform with next generation sequencing readouts. Olink data was analyzed using NPX software,Olink Insights Stat Analysis online tool (<https://olinkproteomics.shinyapps.io/OlinkInsightsStatAnalysis/>), and OlinkAnalyze package using Rstudio Version 1.3. DEPs are provided in **Table S4**. Data are deposited on GEO database under accession #GSE182302.

**Histopathological samples:** Archival tissue from Skin biopsies for diagnostic purposes submitted to the pathology laboratory of dermatopathologist (AD) was used for these studies through an institutional review board (IRB)–approved protocol at UMass Chan (H00020503), and clinical features were re-reviewed by a board-certified dermatologist (MR). All samples were de-identified before use in experiments. Skin biopsies from patients with the diagnosis of ‘skin lupus’ including 5 DLE and 4 SCLE obtained between 2005-2013 were chosen. Three control tissues from healthy margins of skin cancer excisions were also selected from the biorepository from age and sex matched subjects. One DLE and one healthy margin sample were unable to be assessed because of being cut-off from the visualization region on the DSP machine, as well as one SCLE sample due to poor staining, with final n’s of 4 DLE biopsies, 4 SCLE biopsies from 3 patients, and 2 healthy margins. Inclusion criteria for CLE samples included interface dermatitis with perivascular lymphocytic infiltrate, increased dermal mucin on histomorphology and clinical findings consistent with CLE. These patient samples are noted in **Table S2** and in the appropriate results sections.

**Validation dataset analysis for DLE keratinocytes and immune cells:** A separate DSP project was conducted by our collaborators at Yale University School of Medicine. Staining of archived, de-identified human formalin-fixed paraffin-embedded (FFPE) tissues was approved by the Yale University Institutional Review Board (Human Investigative Committee no. 15010105235). This project used the cancer transcriptome atlas (CTA), which includes 1,800+ mRNA probe-based gene targets that cover important aspects of immune response, tumor biology and the microenvironment. We performed comparative analyses between this dataset, comprised of n=3 DLE and n=3 healthy control archival skin tissues, and our DLE keratinocyte and CD3 enriched ROI datasets as a validation cohort using BioVenn *(64)* to visualize overlapping DEGs. These patient samples are noted in **Table S3** and in the appropriate results sections.

**Chemotaxis experiments:** The migration of healthy or lupus donor peripheral blood mononuclear cells (PBMCs) was measured with a HTS Transwell® 96-well Permeable chemotaxis plate (Life science, Coring, Arizona, United states) with 5.0-μm pore filters. In brief, PBMCs were obtained from heparinized blood samples using SepMate™ PBMC isolation tubes (STEMCELL Technologies Inc, Massachusetts), according to the manufacturer's instructions, and were suspended in endotoxin-free RPMI 1640 containing L-glutamine, 5nM HEPES (Corning, Arizona, United States) and 5% Fetal bovine serum (FBS; Sigma Heat Inactivated, US/HI origin). PBMCs were cultured overnight in chemotaxis media containing RPMI and 1% ultrapure BSA prior to use in assays.

The bottom wells were loaded with 200μl of chemotaxis media. We used purified CCL8 (50 ng/ml) CXCL9 (100 ng/ml), CXCL11 (200 ng/ml; all from Biolegend) or CXCL6 (50 ng/ml, R&D Systems), in bottom wells of 5.0-μm plate for chemotaxis of lymphocytes. Chemotaxis medium alone was used as a negative control. The top wells were loaded with 70 μl of PBMC with 1 × 10^5^ cells, and plates were incubated at 37℃ for 2 h. Donors were assayed in triplicate or quadruplicate, and the average value per donor is indicated by a point. Wells were excluded if a bubble was present that precluded migration. The migrated cells in the bottom chamber of each well were collected and stained with CD3, CD8, CD4, CD14, CD16, CD19, CD56, HLA-DR and Live/dead. The number of each type of cells was normalized to count bright beads (Thermo Fisher) and assessed with flow cytometry (Cytek Aurora). The experiment was performed in triplicate or quadruplicate wells with at least 2 donors per chemokine/condition. In tandem, we performed flow cytometry on the input cells using a chemokine receptor staining panel. Blood donors used for chemotaxis assays and chemokine receptor staining are noted in **Table S5** and in the appropriate results sections. Antibody information is provided in **Table S6** and chemokine ligand information is provided in **Table S7**.

**Sample Preparation and Digital Spatial Profiling (DSP):** The NanoString DSP technology provides spatial transcriptomics data using a combination of morphology and immunohistochemical staining*(65, 66)*. Tissue sections at 5 µm thickness were placed on Superfrost Plus slides. Slides were deparaffinized and rehydrated by incubating for 3 × 5 min in CitriSolv, 2 × 5 min in 100% ethanol, 2 × 5 min in 95% ethanol, and 2 × 5 min in deionized water. In order to perform antigen retrieval processing, the slides were put in 1X Citrate Buffer (pH 6) into a pressure cooker at high temperature and high pressure for 15 min. After washing in five times in 1X TBS-T, blocking was performed by putting slides in humidity chambers and covering them with Buffer W (NanoString) for 1 h.

Slides were incubated overnight at 4°C overnight in a humidity chamber with 1.25 ug/ml anti-SMA, 5 ug/ml anti-CD45, and 1 ug/ml anti-CD3 antibodies. Slides were washed three times in 1X TBS-T post-staining and then post-fixed in 4% PFA. After two additional washes in 1X TBS-T to remove the fixative, nuclei were stained with 500 nM SYTO83 for 15 min at room temperature followed by one wash in 1X TBS-T before loading onto the GeoMx instrument.

Regions of interest (ROI) using the polygon selection tool were created and cell masking was performed using the morphology markers to define cell type as follows: CD45+CD3+ for T cells, CD45+CD3- for non-T cell-immune cells, CD45- and morphology for keratinocytes, CD45-SMA+ for endothelial cells.

For Whole Transcriptome Atlas (WTA), photocleaved oligos were then collected for sequencing. For protein profiling, we used 62 immune related biomarkers with the following panels: Immune Cell Profiling (18 markers), immuno-oncology (IO) Drug Target (10 markers), Immune Activation Status (8 markers), Immune Cell Typing (7 markers), Cell Death (10 markers), and PI3K/AKT Signaling (9 markers).

### **Library preparation and sequencing for WTA**: Oligos from each ROI were collected via microcapillary tube inspiration using the DSP platform robotic system and transferred into a microwell plate for use in whole transcriptome atlas (WTA) sequencing, which includes 18,000+ protein encoding mRNA targets that encompasses the whole human transcriptome. Collected oligos were amplified using a forward primer and a reverse primer that serve as Illumina i5/i7 unique dual indexing sequences to index ROI identity. After purifying the PCR products with AMPure XP beads (Beckman Coulter), they were sequenced. Library purity and concentration were measured with DNA Bioanalyzer chip (Agilent). Data are deposited on GEO Database under accession # GSE182825.

**nCounter Readout for Protein DSP**: Protein modules were collected and run in the MAX machine at the Comparative Pathobiology and Genomics Shared Resources (“CPGSR”) in Cummings School of Veterinary Medicine at Tufts University.

**Data processing and analysis:** Reads after sequencing, were trimmed, merged, and aligned to retrieve the identity of probes. PCR duplicates and duplicate reads were removed and the reads were converted to digital counts. The RNA sequencing saturation was sufficient and above 50%. After removing the outlier probes, the mean of the individual probe counts is considered as the reported count value. using GeoMX software (NanoString), 75% upper quartile (Q3) of the counts per ROI were selected after removing genes with zero counts. The Q3 normalized counts were compared across ROI and disease subtypes using several approaches. Overall differences were determined using linear mixed model analysis in GeoMX software. QC of data was performed using GeoMX software, and normalized counts were uploaded to ROSALIND software for comparison (as in Fig S1, <https://rosalind.bio/>, *(67)*). Specific groups of genes of interest, including chemokines/chemokine receptors, were analyzed as groups using GraphPad Prism version 9 with two-way ANOVA and Tukey’s post hoc tests. Median values for groups and ROIs generated in GeoMX software were used to visualize large gene sets as heatmaps using GraphPad Prism.

**Reanalysis of GSE112943 dataset:** Microarray analysis of bulk RNA was previously performed on matched biobanked samples*(18)*. To compare bulk RNA from spatial transcriptomics, we used Geo2R to download the toptable of significant genes for CCLE/DLE versus healthy and SCLE versus healthy, and compared DEGs with P<0.01 to DEGs from all ROIs pooled from the dataset presented in this manuscript using BioVenn *(64)*.

**Data visualization tools:** Gene Ontology (GO) analysis*(68)*, Gene Set Enrichment Analysis (GSEA)*(69)* were performed using GeoMX software. The CellPhone Database*(70)* was queried for cell-cell communication pathway analyses in different ROI types. Venn diagrams were generated with BioVenn software*(64)*, and hierarchical clustering were performed with ClustVis*(71)* and/or Morpheus software (<https://software.broadinstitute.org/morpheus>).

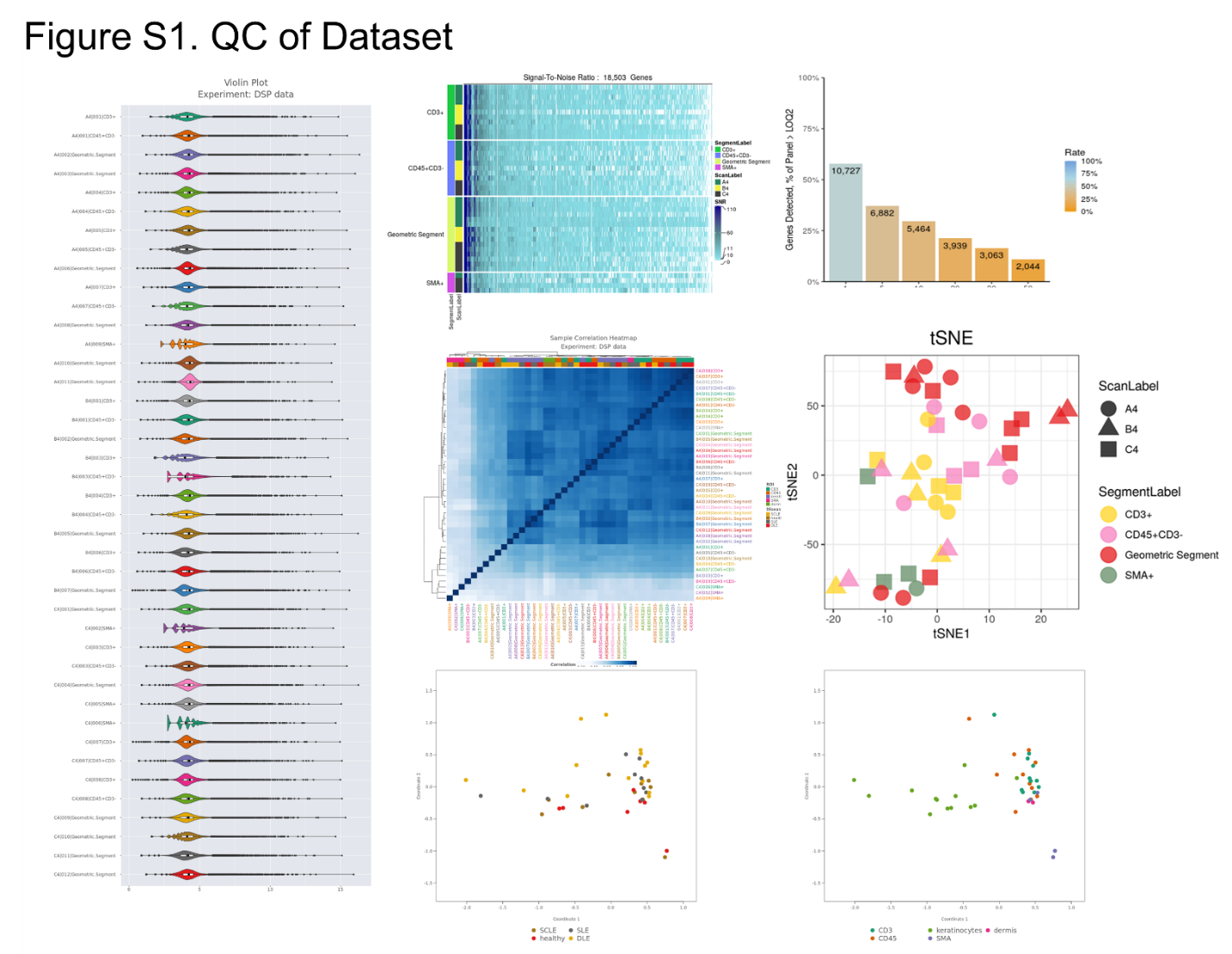

**Fig. S1.** **Quality Control for Digital Spatial Profiling (DSP) Whole Transcriptome Atlas (WTA) spatial transcriptomics**. (A). Violin Plot showing sample distribution. (B). Signal-to-noise heatmap grouped by ROI type and slide. (C). Histogram Chart for LOQ2 Values, presents a histogram chart that conveys the distribution of LOQ2 values. These values were employed to filter genes that are expressed near background levels, aiding in data quality assessment. (D). Sample Correlation Heatmap showed correlation heatmap is presented, depicting the pairwise correlation between samples within the DSP - RNA dataset. Notably, this heatmap highlights specific regions of interest (ROIs) in proximity to the respective antibodies used for staining. (E). t-distributed Stochastic Neighbor Embedding (t-SNE) Graph visualizes the data points with scan labels and segment labels, effectively illustrating the spatial distribution of data and the relationship between segments. (F). Principal Component Analysis (PCA) for the clustering of samples based on disease type (left) and cell types (right).

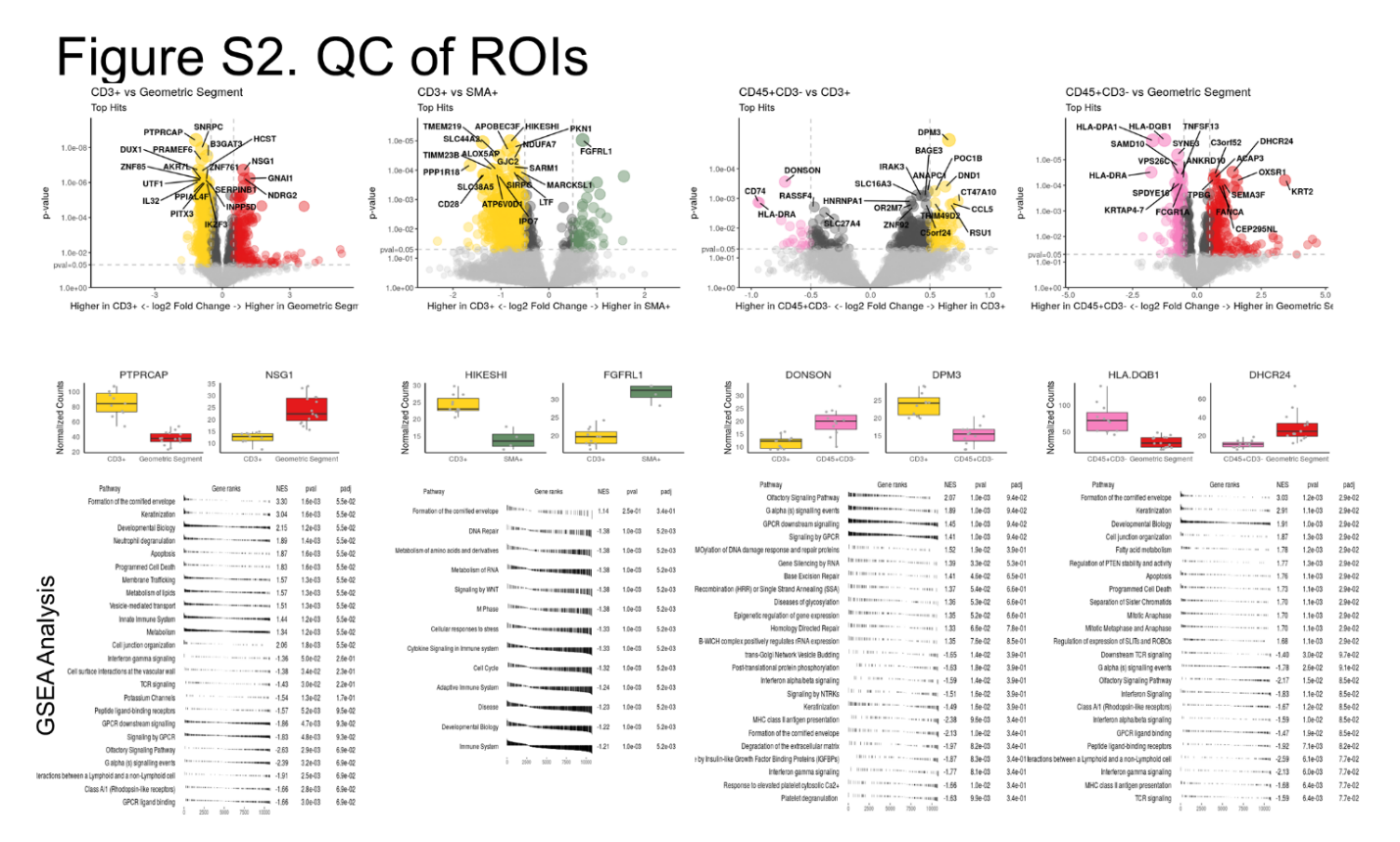

**Fig S2.** **Quality Control and Pathway Analysis for Regions of Interest (ROIs)**. (A). Volcano Plot Comparisons present a comparing gene expression in different segments. The following pairwise segment comparisons are CD3+ vs. Geometric, CD3+ vs. SMA+, CD45+CD3- vs. CD3+, and CD45+CD3- vs. Geometric. Each volcano plot illustrates the distribution of gene expression across segments, with the log-fold change on the x-axis and the statistical significance on the y-axis. Additionally, for each volcano plot, a bar chart highlights the most highly expressed genes in the respective segments, providing valuable insights into the key drivers of variation. (B). Gene Set Enrichment and Analysis (GSEA) unveils significant pathways and functional annotations associated with the gene expression data within the ROIs.

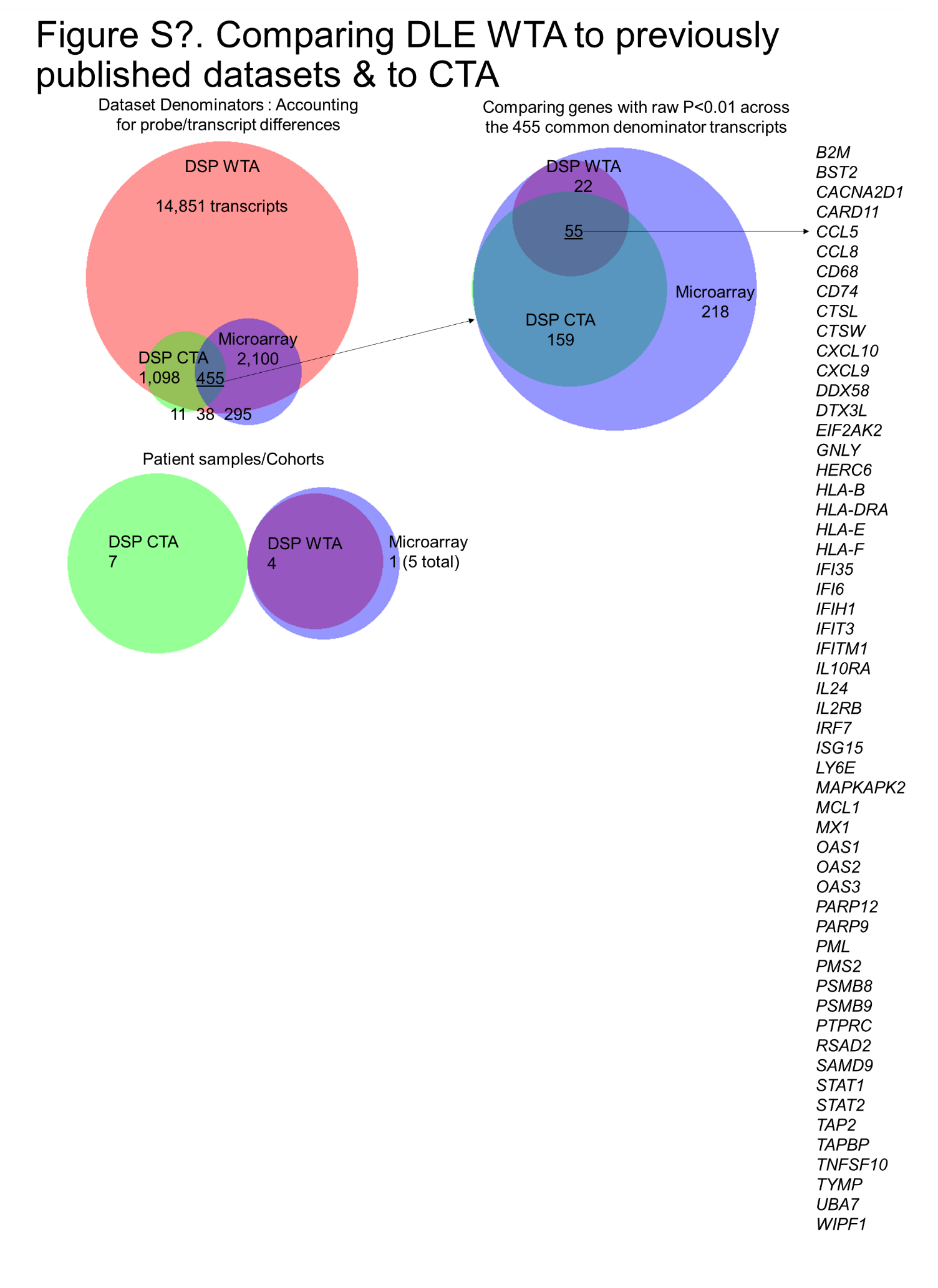

**Fig S3. Validation of DSP dataset using historical microarray dataset and a cancer transcriptome atlas (CTA) dataset.** We compared DEGs in DSP WTA (this dataset), DSP CTA (Vesely dataset) and microarray (Ko/Harris dataset). Among these datasets, 455 common probe/transcript denominators were identified. This figure focuses on the 55 genes present across all datasets with raw p-values < 0.01. This comparative analysis offers a robust assessment of the consistency and reliability of the DSP WTA dataset by examining the shared DEGs across multiple datasets. The presence of these 55 genes underscores their significance in the context of cutaneous lupus.

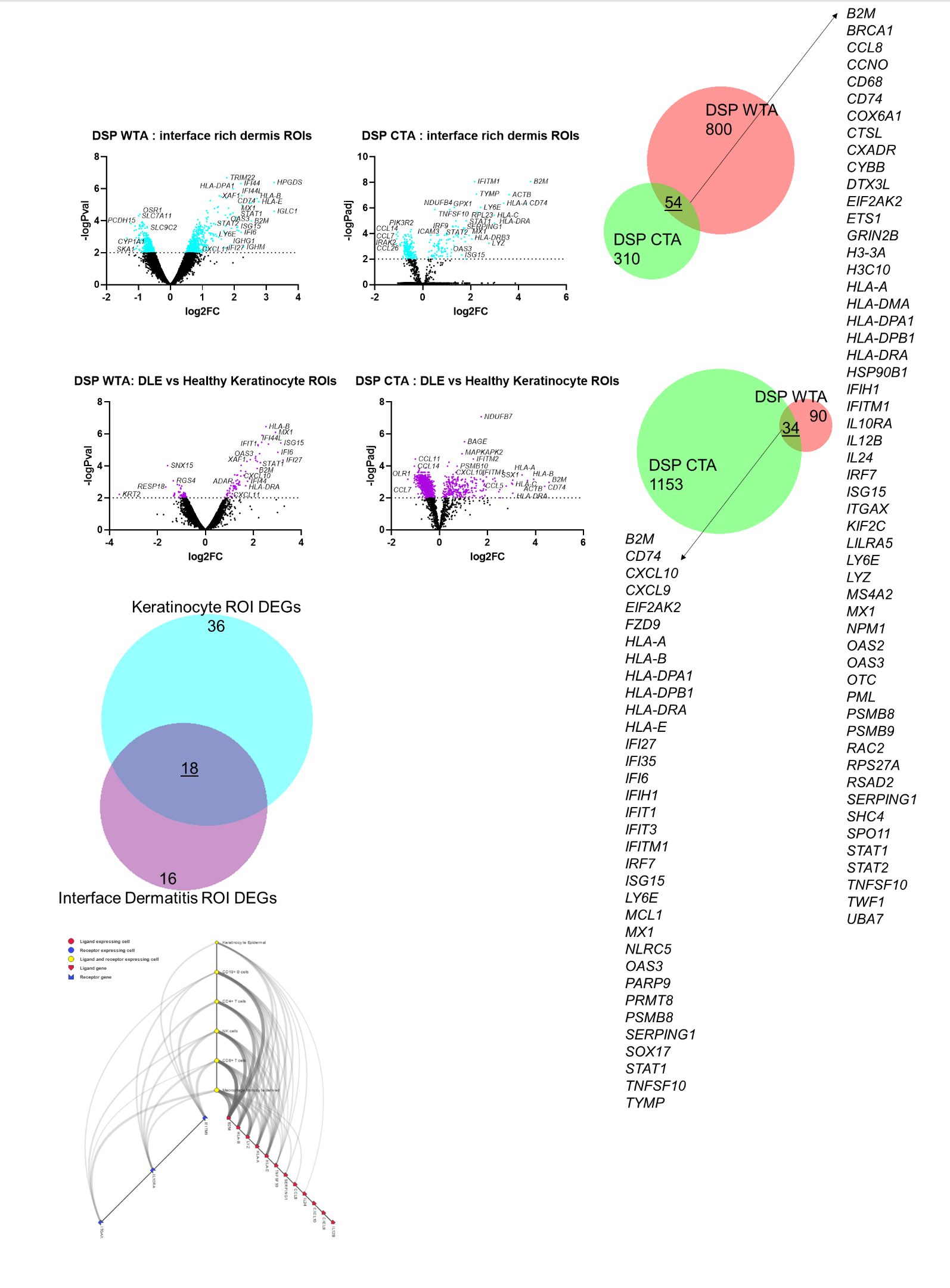

**Fig S4. Validation of DLE dataset ROIs as compared to CTA.** We compared DEGs in interface dermatitis rich ROIs (CD45+) in WTA (this dataset) and CTA (Vesely dataset) (teal volcanoes). Of these, we found 54 conserved DEGs in the inflammatory infiltrate. We also compared DEGs in the epidermal/keratinocyte geometric ROIs in the WTA and CTA datasets and found 34 overlapping DEGs (purple volcanoes). Last, we examined shared DEGs across ROI types and found 18 DEGs that are expressed by both the stroma and the immune cells (purple vs teal BioVenn). We also examined receptor:ligand pairs across these ROI types to understand how the immune system communicates with keratinocytes in CLE using cellPhoneDB, which demonstrated multiple predicted interactions.

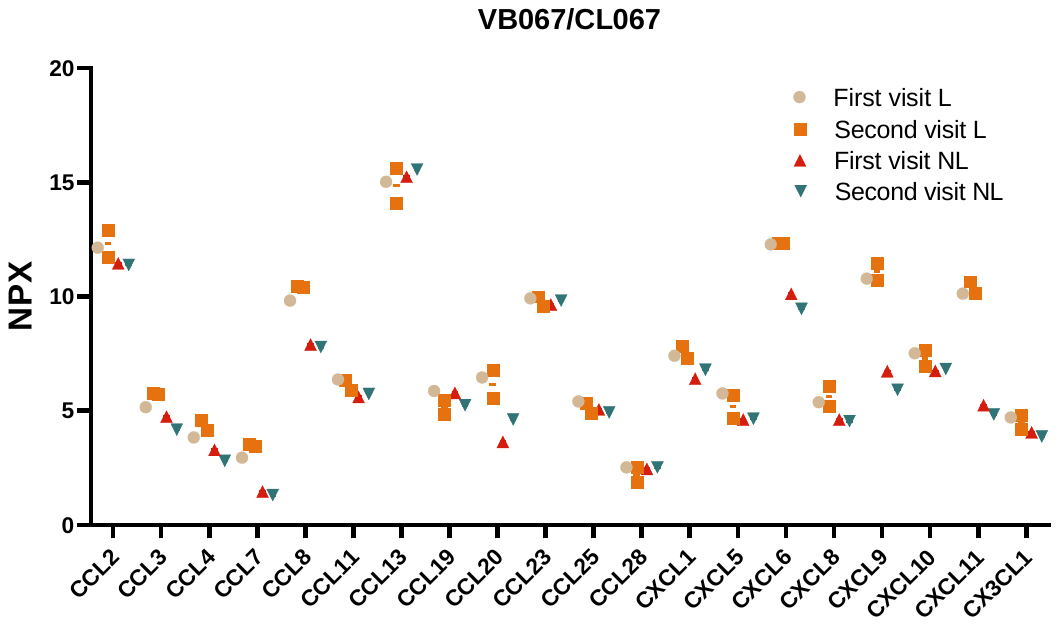

**Fig S5.** **Stability of chemokines in lesional and nonlesional samples over time**. We compared data obtained during the first and second visits from a repeat donor who was sampled approximately 6 months apart. The quantification of chemokines is based on Normalized Protein eXpression (NPX) values from 96-plex immunoassay targeted proteomics.

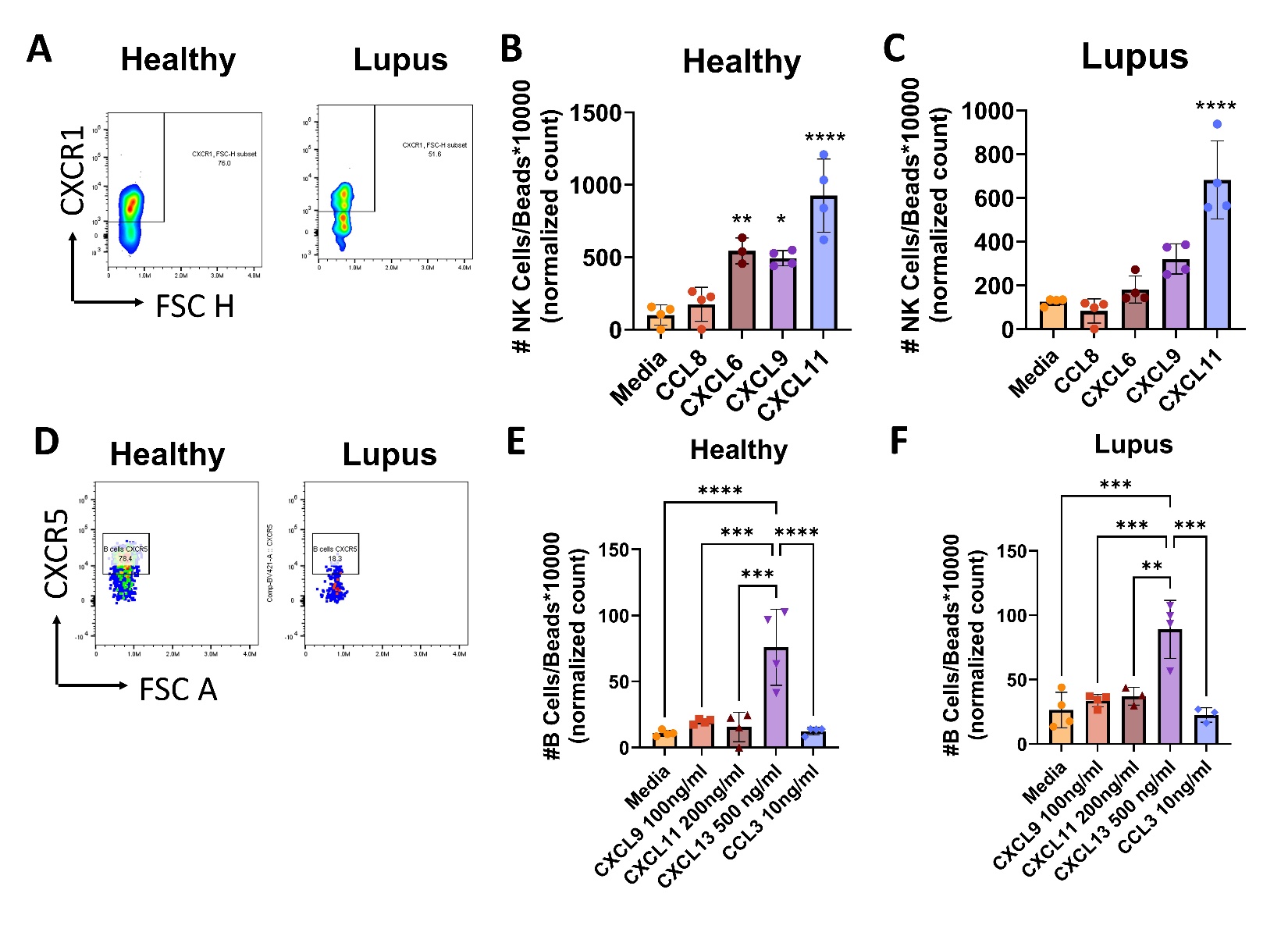

**Fig** **S6. Assessment of NK cell and B cell migration**. We tested n=3 healthy and n=2 lupus donor PBMCs in triplicate or quadruplicate in chemotaxis assays. A. Representative CXCR1 staining on NK cells (pregated on live, single CD45+CD3-CD19-CD56+). Representative chemotaxis assay from a B. healthy donor and C. lupus donor demonstrating migration towards CXCL6<CXCL9<CXCL11. D. Representative CXCR5 staining on B cells (pregated on live, single CD45+CD3-CD19+). Representative chemotaxis assay from a B. healthy donor and C. lupus donor demonstrating migration towards CXCL13. (one-way ANOVAs with significant posttests as indicated).

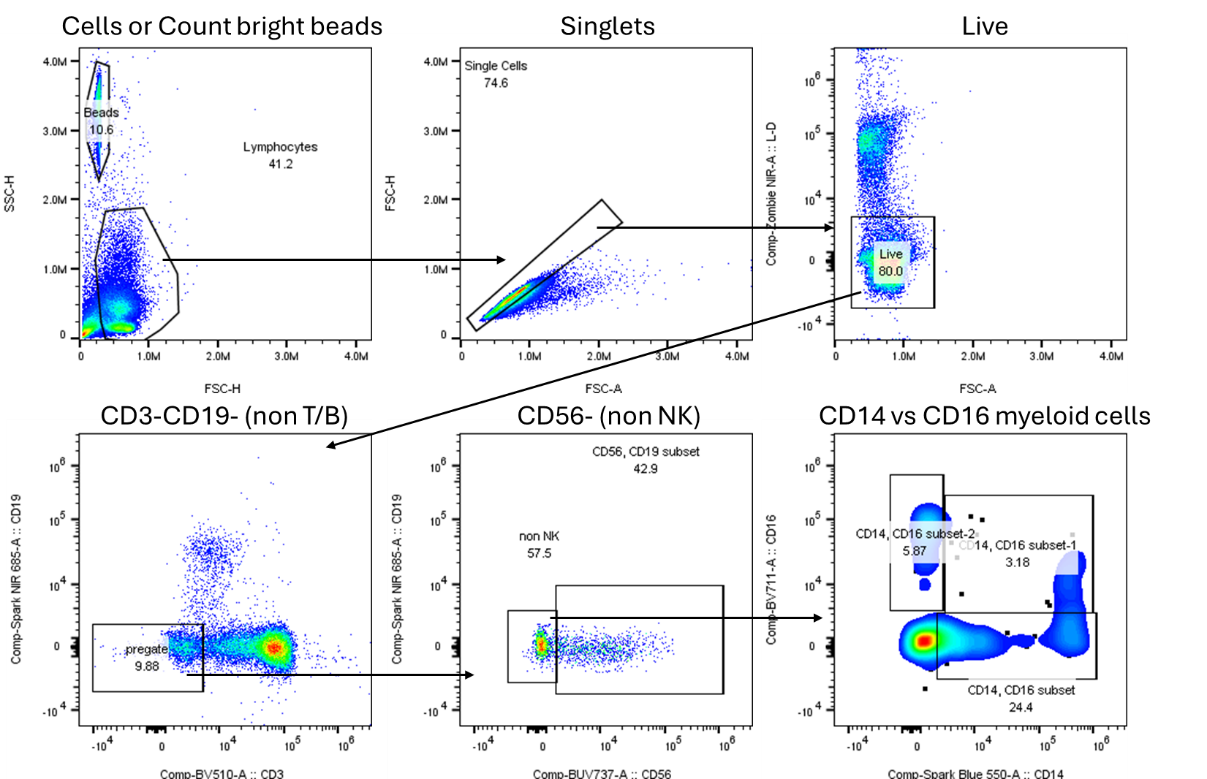

**Fig S7. Flow gating strategy for assessing myeloid populations**. Example flow gating strategy employed to assess CD14 vs CD16 myeloid cells using cells->singlets->live->CD3-CD19- and CD56- pregates, followed by CD14 vs CD16. This example is from PBMCs.

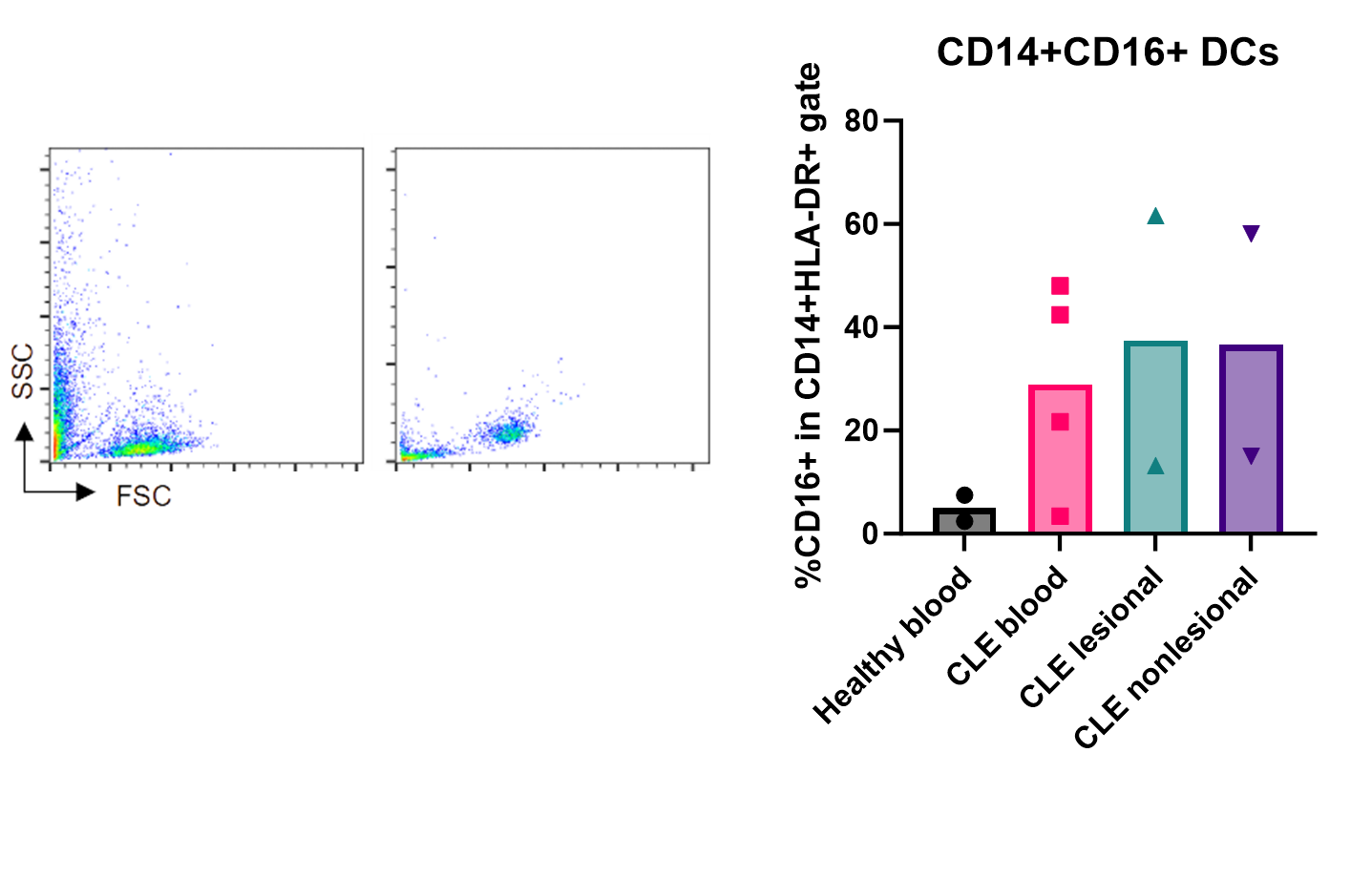

**Fig S8. Examination of CD14+CD16+ cells in blister biopsies**. Confirmation of CD16+ antigen presenting cell populations as described in Kahlenberg 10X spatial dataset. Note that not every blister and blood donor had significant cell populations as detected by flow cytometry.

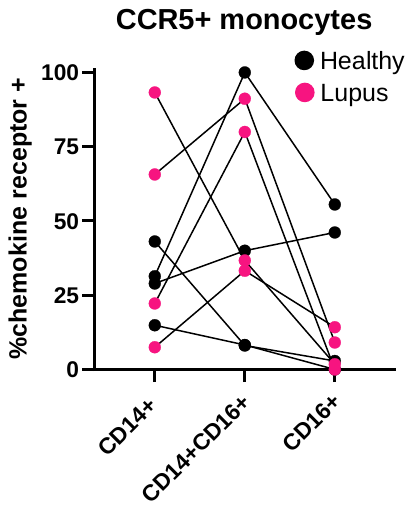

**Fig S9**. **CCR5 is not enriched on CD14 vs CD16 expressing myeloid cells**. CCR5 is an alternate receptor for CCL8 on monocytes. We noted no specific trend in expression in different quadrants for CD14 vs CD16 expressing myeloid cells, which could not account for chemotactic differences observed in response to CCL8.

**Table S1. Blister biopsy and blood donation patient info & characterization (UMass Chan).**

| **Subject #** | **Diagnosis** | **Age** | **Sex** | **Race/Ethnicity** | **Treatment status** | **Pertinent medical history** | **Blister biopsy** | **Blister site L** | **Blister site NL** | **Blood sample** |
| --- | --- | --- | --- | --- | --- | --- | --- | --- | --- | --- |
| 1 | SCLE | 45 | M | White | Hydroxychloroquine (Plaquenil), last used 2 years prior to the biopsy; triamcinolone 0.1% ointment BID, PRN 1 year prior; Steroids Creams Last applied few months prior to the biopsy | none | Y | upper back | forearm, inner aspect | Y |
| 2 | Active SLE | 44 | F | White, Hispanic Latina | On Lyrica | none | Y | cheek | forearm | Y |
| 3 | SCLE | 75 | F | White | Hydroxychloroquine (Plaquenil). Prescribed just prior to biopsy: Protopic 0.1% ointment BiD PRN itching or rash on the face; triamcinolone 0.1% cream BID PRN itching or rash on the trunk and limbs; fluocinonide scalp solution | Raynaud's | Y | outer arm | forearm, inner aspect | Y |
| 4 | SCLE with SLE | 55 | F | White | On hydroxychloroquine | Fibromyalgia, photosensitivity | Y | arm | arm | Y |
| 5 | Healthy | 34 | F | White | none | none | N | N/A | N/A | Y |
| 6 | Healthy | 54 | F | Asian | none | none | N | N/A | N/A | Y |
| 7 | Healthy | 28 | M | White, Middle Eastern | none | none | Y | N/A | arm | N |
| 8 | Healthy | 60 | F | White | none | none | Y | N/A | arm | N |
| 9 | Healthy | 52 | F | White | none | none | Y | N/A | arm | Y |
| 10 | Healthy | 56 | M | ND | Chlorthalidone, Lisinopril, Lovaza, Myrbetriq, Tamsulosin, Omeprazole, gabapentin; Triamcinolone for dry skin on legs | High blood pressure, GERD, dry skin, prostate hyperplasia | Y | N/A | arm | N |
| 11 | Healthy | 26 | F | White | none | none | Y | N/A | arm | N |
| 12 | SCLE | 74 | F | White | Hydroxychloroquine, triamcinolone, methotrexate, vitamin D, paroxetine | Sicca syndrome (HCC) | Y | Right lower arm, left lower arm | Left upper thigh | Y |
| 13 | SCLE | 45 | F | White | Betamethasone, cyclobenzaprine HCL, diclofenac sodium, tolterodine tartrate, escitalopram oxalate | none | Y | Left upper arm | Left lateral abdomen | Y |

SCLE = subacute cutaneous lupus erythematosus; SLE = systemic lupus erythematosus; BID = twice daily; PRN = taken as needed; FH = family history; L = lesional; NL = nonlesional; N/A = not applicable, ND = not determined/disclosed, Y = yes, N = no

**Table S2. Archival CLE biopsies used in WTA Digital Spatial Profiling (UMass Chan).**

| **Disease status** | **Systemic symptoms** | **Age (yr)** | **Sex** | **Ethnicity** | **Age at diagnosis (yr)** | **Age at skin biopsy (yr)** | **Site of Bx** | **Positive antibodies** | **CLE-related medications at the time of Bx** |
| --- | --- | --- | --- | --- | --- | --- | --- | --- | --- |
| SCLE | Bilateral chronic hand pain | 68 | M | White, not Hispanic Latino | 68 | 68 | Shoulder+ Forearm | +ANA (speckled), +SSA, +dsDNA | None |
| DLE | Swollen salivary glands | 42 | M | White, not Hispanic Latino | ? | 42 | Face | +Anti-SM/RNP, +ANA | HCTZ (new forearm rash). Cheek rash predated HCTZ |
| SCLE (drug induced) | None | 76 | F | White, not Hispanic Latino | 76 | 76 | Shoulder | +ANA (speckled), +SSA, +SSB; Later diagnosed with Sjogren's | HCTZ, glipizide, omeprazole |
| SCLE | None (until 04/2020, joint pain and blue toes) | 37 | M | White, not Hispanic Latino | 37 | 37 | Abdomen | +ANA, +SSA, +SSB | Oral ketoconazole (previous hx of tinea versicolor) |
| DLE | None | 68 | F | White, not Hispanic Latino | 68 | 68 | Scalp | +ANA (speckled and homogeneous),+SSA, +Sm/RNP | Topical clobetasol, T-Gel shampoo (initially thought psoriasis) |
| DLE | None | 45 | F | White, not Hispanic Latino | 45 | ? | Neck (Posterior auricular region) | Borderline +ANA | ? |
| DLE | vague arthralgia, no other features to meet criteria for SLE | 51 | F | White, not Hispanic Latino | 51 | 51 | Forehead | +ANA, +dsDNA | None |
| Healthy | N/A | 64 | F | White, not Hispanic Latino | N/A | N/A | Forehead | N/A | None |
| Healthy | N/A | 32 | M | White, not Hispanic Latino | N/A | N/A | Forehead | N/A | None |
| Healthy | N/A | 41 | M | White, not Hispanic Latino | N/A | N/A | Forearm | N/A | None |

**Table S3. Validation archival CLE biopsies used in CTA Digital Spatial Profiling (Yale).**

| **Sample Name** | **Sex** | **Age (years)** | **Location** | **Treatment at time of biopsy** | **ROIs** |
| --- | --- | --- | --- | --- | --- |
| ***CTA panel*** |  |  |  |  |  |
| DLE #1 | F | 45 | Cheek | None | 1-4 |
| DLE #2 | M | 53 | Nose | None | 5-8 |
| DLE #3 | F | 40 | Cheek | None | 9-12 |
| Control #1 | F | 28 | Neck | - | 1-4 |
| Control #2 | M | 52 | Back | - | 5-8 |
| Control #3 | M | 33 | Back | - | 9-12 |

**Table S4. 96-plex immunoassay DEPs calculated by NPX software and 2-way ANOVA.**

| Significant for CLE lesional versus nonlesional and healthy | | | | | |  | |  | |
| --- | --- | --- | --- | --- | --- | --- | --- | --- | --- |
| CXCL11 | |  |  |  | |  | |  | |
| CLE Lesional vs. CLE Nonlesional | | 2.878 | 0.2467 to 5.508 | Yes | | * | | 0.0351 | |
| CLE Lesional vs. Healthy | | 5.395 | 1.531 to 9.259 | Yes | | * | | 0.0127 | |
| CLE Nonlesional vs. Healthy | | 2.518 | -2.698 to 7.733 | No | | ns | | 0.1869 | |
| CXCL9 | |  |  |  | |  | |  | |
| CLE Lesional vs. CLE Nonlesional | | 2.74 | 1.189 to 4.291 | Yes | | ** | | 0.0023 | |
| CLE Lesional vs. Healthy | | 5.385 | 0.2335 to 10.54 | Yes | | * | | 0.0456 | |
| CLE Nonlesional vs. Healthy | | 2.644 | -1.693 to 6.981 | No | | ns | | 0.167 | |
| CXCL6 | |  |  |  | |  | |  | |
| CLE Lesional vs. CLE Nonlesional | | 1.97 | 0.2310 to 3.710 | Yes | | * | | 0.0275 | |
| CLE Lesional vs. Healthy | | 5.917 | 0.08853 to 11.75 | Yes | | * | | 0.0482 | |
| CLE Nonlesional vs. Healthy | | 3.947 | -2.539 to 10.43 | No | | ns | | 0.137 | |
| IFN-gamma | |  |  |  | |  | |  | |
| CLE Lesional vs. CLE Nonlesional | | 2.358 | 0.2003 to 4.516 | Yes | | * | | 0.0331 | |
| CLE Lesional vs. Healthy | | 4.2 | 1.447 to 6.953 | Yes | | ** | | 0.0075 | |
| CLE Nonlesional vs. Healthy | | 1.842 | -0.8639 to 4.548 | No | | ns | | 0.1426 | |
| CASP-8 | |  |  |  | |  | |  | |
| CLE Lesional vs. CLE Nonlesional | | 1.746 | 0.1036 to 3.389 | Yes | | * | | 0.0374 | |
| CLE Lesional vs. Healthy | | 3.206 | 0.6071 to 5.804 | Yes | | * | | 0.0235 | |
| CLE Nonlesional vs. Healthy | | 1.459 | -1.249 to 4.168 | No | | ns | | 0.2286 | |
| CTF1 | |  |  |  | |  | |  | |
| CLE Lesional vs. CLE Nonlesional | | 0.7969 | 0.08803 to 1.506 | Yes | | * | | 0.0287 | |
| CLE Lesional vs. Healthy | | 1.008 | 0.2062 to 1.810 | Yes | | * | | 0.0177 | |
| CLE Nonlesional vs. Healthy | | 0.2112 | -0.4943 to 0.9167 | No | | ns | | 0.6099 | |
| Significant for CLE lesional or nonlesional versus healthy | | | | |  | |  | |  |
| HGF |  | |  | |  | |  | |  |
| CLE Lesional vs. CLE Nonlesional | 1.419 | | -0.3617 to 3.200 | | No | | ns | | 0.1258 |
| CLE Lesional vs. Healthy | 4.07 | | 2.410 to 5.730 | | Yes | | *** | | 0.0004 |
| CLE Nonlesional vs. Healthy | 2.651 | | 0.8788 to 4.423 | | Yes | | ** | | 0.007 |
| Flt3L |  | |  | |  | |  | |  |
| CLE Lesional vs. CLE Nonlesional | 0.005873 | | -0.7783 to 0.7900 | | No | | ns | | 0.9998 |
| CLE Lesional vs. Healthy | 2.871 | | 0.6221 to 5.121 | | Yes | | * | | 0.0265 |
| CLE Nonlesional vs. Healthy | 2.865 | | 0.5416 to 5.189 | | Yes | | * | | 0.0299 |
| CCL28 |  | |  | |  | |  | |  |
| CLE Lesional vs. CLE Nonlesional | 0.1427 | | -0.3112 to 0.5966 | | No | | ns | | 0.687 |
| CLE Lesional vs. Healthy | 0.5747 | | 0.1965 to 0.9529 | | Yes | | ** | | 0.0063 |
| CLE Nonlesional vs. Healthy | 0.432 | | 0.04441 to 0.8196 | | Yes | | * | | 0.0311 |
| CCL25 |  | |  | |  | |  | |  |
| CLE Lesional vs. CLE Nonlesional | -0.1353 | | -1.333 to 1.062 | | No | | ns | | 0.951 |
| CLE Lesional vs. Healthy | 2.122 | | 0.3620 to 3.881 | | Yes | | * | | 0.027 |
| CLE Nonlesional vs. Healthy | 2.257 | | 0.5139 to 4.000 | | Yes | | * | | 0.0192 |
| IFNL1 |  | |  | |  | |  | |  |
| CLE Lesional vs. CLE Nonlesional | 1.479 | | -0.5759 to 3.534 | | No | | ns | | 0.1587 |
| CLE Lesional vs. Healthy | 2.646 | | 0.5894 to 4.703 | | Yes | | * | | 0.0164 |
| CLE Nonlesional vs. Healthy | 1.167 | | 0.2188 to 2.116 | | Yes | | * | | 0.0203 |
| CEACAM3 |  | |  | |  | |  | |  |
| CLE Lesional vs. CLE Nonlesional | 0.1451 | | -0.02555 to 0.3157 | | No | | ns | | 0.096 |
| CLE Lesional vs. Healthy | -0.296 | | -0.5794 to -0.01256 | | Yes | | * | | 0.043 |
| CLE Nonlesional vs. Healthy | -0.4411 | | -0.7595 to -0.1226 | | Yes | | * | | 0.0218 |

**Table S5. Chemotaxis and chemokine receptor staining blood donor information (UMass Chan and Dartmouth Hitchcock).**

| **Subject** | **Gender** | **Age** | **race** | **Skin color** | **Diagnosis** | **Treatment** | **Chemotaxis ligands** | **Chemokine Receptor staining** |
| --- | --- | --- | --- | --- | --- | --- | --- | --- |
| A | Female | 32 y/o | Hispanic | white | SLE | Plaquenil 300mg daily | N/A | CCR1, CXCR3, CCR5, CCR7, CXCR1, CXCR5, CXCR2, CCR2 |
| B | Female | 64 y/o | Hispanic | white | DLE/SCLE | 6 (15mg) pills MTX/week for past 8 mo, synthroid, albuterol | CXCL9, 11,13, CCL3 | CCR1, CXCR3, CCR5, CCR7, CXCR1, CXCR5, CXCR2, CCR2 |
| C | Male | 42 y/o | Hispanic | White | DLE with SLE | Benlysta 200mg Qweekly, methotrexate 25mg Qweekly, plaquenil 200mg, Kenalog, folic acid 1mg QD, betamethasone | CXCL9, 11, 13, 6, CCL3, 25, 8 | CCR1, CXCR3, CCR5, CCR7, CXCR1, CXCR5, CCR9, CCR2 |
| D | Female | 43 y/o | White | White | CLE/Sjorgens | Plaquenil | N/A | CCR1, CXCR3, CCR5, CCR7, CXCR1, CXCR5, CXCR2, CCR2 |
| E | Female | 48  y/o | White | White | CCLE with SLE, also has non scarring alopecia | Triamcinolone 0.1% PRN, protopic 0.1% PRN, hydroxychloroquine 300mg QD | N/A | CCR1, CXCR3, CCR5, CCR7, CXCR1, CXCR5, CCR9, CCR2 |
| F | Female | 35 y/o | Middle eastern | White | Healthy |  | N/A | CCR1, CXCR3, CCR5, CCR7, CXCR1, CXCR5, CXCR2, CCR2 |
| G | Female | 34 y/o | Middle eastern | White | Healthy |  | CXCL9, 11,13, CCL3 | CCR1, CXCR3, CCR5, CCR7, CXCR1, CXCR5, CXCR2, CCR2 |
| H | Female | 36 y/o | Hispanic | White | Healthy |  | CXCL9, 11, 13, 6, CCL3, 25, 8 | CCR1, CXCR3, CCR5, CCR7, CXCR1, CXCR5, CCR9, CCR2 |
| I | Male | 28 y/o | Middle eastern | White | Healthy; Hypothyroidism | Levothyroxine | CXCL9, 11, 6, CCL3, 8 | CCR1, CXCR3, CCR5, CCR7, CXCR1, CXCR5, CCR9, CCR2 |
| J | Female | 28 y/o | Middle eastern | White | Healthy |  | CXCL9, 11, 13, 6, CCL3, 25, 8 | CCR1, CXCR3, CCR5, CCR7, CXCR1, CXCR5, CCR9, CCR2 |
| K | Male | 26 y/o | Asian | White | Healthy |  | CXCL6 dose curve | CCR1, CXCR3, CCR5, CCR7, CXCR1, CXCR5, CCR9, CCR2 |
| L | Male | 43 y/o | Hispanic | White | CLE with SLE | Hydroxychloroquine, methylprednisolone acetate, mometasone, prednisone, triamcinalone | CXCL9, 11, 6, CCL3, 8 | CCR1, CXCR3, CCR5, CCR7, CXCR1, CXCR5, CCR9, CCR2 |

DLE = discoid lupus erythematosus; SCLE = subacute cutaneous lupus erythematosus; SLE = systemic lupus erythematosus; N/A = not applicable

| M | Female | 26 y/o |  | White | DLE | Vit D, HCQ 450 mg Daily, Benlysta 200 mg, Triamcinolone | CXCL6, CCL8 | CCR1, CXCR3, CCR5, CCR7, CXCR1, CXCR5, CCR9, CCR2 |
| --- | --- | --- | --- | --- | --- | --- | --- | --- |
| N | Male | 43 y/o | Hispanic or Latino | White | SLE, from medical note: "Erythematous burning dermatitis on chest; from reviewing pictures I am not convinced that this was related to cutaneous lupus (definitely not consistent with discoid lupus, possible it was subacute cutaeous lupus but the lack of epidermal changes makes it less likely)" | Predinosone, Hydroxychloroquine | CXCL9, 11, 6, CCL3, 8 | CCR1, CXCR3, CCR5, CCR7, CXCR1, CXCR5, CCR9, CCR2 |
| O | Female | 49 y/o | Black or African American | Not Hispanic or Latino | DLE, SLE, Lupus Alopecia | Hydroxychloroquine, Carboxymethylcellulose sodium 0.5% eye drops, (Unknown if she is taking meds for hypertension) | CXCL6, CCL8 | CCR1, CXCR3, CCR5, CCR7, CXCR1, CXCR5, CCR9, CCR2 |
| P | Male | 57 y/o | Asian | Not Hispanic or Latino | Healthy | Metformin (T2 Diabetes), Atorvastatin (Hyperlipidemia), Hyzaar (Hypertension) | CXCL6, CCL8 | CCR1, CXCR3, CCR5, CCR7, CXCR1, CXCR5, CCR9, CCR2 |
| Q | Female | 40y/o | Prefer not to answer | Hispanic or Latino | DLE/Tumid Lupus, SLE | Hydroxychloroquine (DLE/SLE), Trulicity (Prediabetes), Cevimeline (Sjogrens) | CXCL6, CCL8 | CCR1, CXCR3, CCR5, CCR7, CXCR1, CXCR5, CCR9, CCR2 |
| R | Female | 26 y/o | White | Hispanic or Latino | Healthy | Baclofen | CXCL6, CCL8 | CCR1, CXCR3, CCR5, CCR7, CXCR1, CXCR5, CCR9, CCR2 |
| S | Female | 35 y/o | White | Not Hispanic or Latino | Healthy | Albuterol inhaler (Asthma), Mirena | CXCL6, CCL8 | CCR1, CXCR3, CCR5, CCR7, CXCR1, CXCR5, CCR9, CCR2 |

DLE = discoid lupus erythematosus; SCLE = subacute cutaneous lupus erythematosus; SLE = systemic lupus erythematosus; N/A = not applicable

**Table S6. Antibody information & RRIDs.**

| **Marker** | **Company** | **Catalog No.** | **Lot No.** | **Dose** | **RRID** |
| --- | --- | --- | --- | --- | --- |
| CD3 (BV510) | Biolegend | 344828 | B370758 | 5ul/test | RRID:AB_2563704 |
| CD8 (BV605) | Biolegend | 344742 | B370756 | 5ul/test | RRID:AB_2566513 |
| CD4 (Alexa Flour 700) | Biolegend | 344622 | B365788 | 5ul/test | RRID:AB_2563150 |
| CD14 (Spark Blue 550) | Biolegend | 367148 | B361330 | 5ul/test | RRID:AB_2832724 |
| CD56 (BUV737) | BD Biosciences | 612766 | 2332923 | 5ul/test | RRID:AB_2813880 |
| CD19 (Spark Nir 685) | Biolegend | 302270 | B358744 | 5ul/test | RRID:AB_2832581 |
| CD16 (BV711) | Biolegend | 360732 | B397395 | 5ul/test | RRID:AB_2800992 |
| HLA-DR (PE) | Biolegend | 307606 | B373145 | 5ul/test | RRID:AB_314684 |
| CCR1 (PerCP/Cyanine 5.5) | Biolegend | 362912 | B323489 | 5ul/test | RRID:AB_2728353 |
| CXCR3 (PECy7) | Biolegend | 353720 | B322685 | 5ul/test | RRID:AB_11219383 |
| CCR5 (BUV563) | BD Biosciences | 741401 | 3198452 | 5ul/test | RRID:AB_2870893 |
| CCR7 (PE/Fire 640) | Biolegend | 353262 | B351674 | 5ul/test | RRID:AB_2876669 |
| CXCR1 (APC) | Biolegend | 320612 | B397637 | 5ul/test | RRID:AB_2126475 |
| CXCR5 (BV421) | Biolegend | 356920 | B381006 | 5ul/test | RRID:AB_2562303 |
| CXCR2 (PE/Dazzle 594) | Biolegend | 320722 | B381595 | 5ul/test | RRID:AB_2750215 |
| CCR2 (BV785) | Biolegend | 357234 | B366953 | 5ul/test | RRID:AB_2800972 |
| CCR9 (PE/Dazzle 594) | Biolegend | 358918 | B384511 | 5ul/test | RRID:AB_2715935 |
| Live/Dead (Zombie NIR Dye) | Biolegend | 77184 | B369131 | 5ul/test | N/A |

**Table S7. Chemokine information.**

| **Reagent** | **Company** | **Catalog #** | **Lot #** | **Dose (s) tested in chemotaxis assay** |
| --- | --- | --- | --- | --- |
| CCL3 | Biolegend | 759504 | B364645 | 10, 50ng/mL |
| CCL19 | Biolegend | 582104 | B361963 | 30 ng/ml |
| CCL23 | Biolegend | 587002 | B389072 | 10 ng/ml |
| CCL25 | Biolegend | 586804 | B272484 | 200 ng/ml |
| CXCL6 | R&D Systems | 333-GC-025/CF | AMM0222041 | 5, 50, 100, 200, 500 ng/ml |
| CXCL9 | Biolegend | 578104 | B366484 | 100 ng/ml |
| CXCL11 | Biolegend | 574904 | B312963 | 200ng/mL |
| CXCL13 | Biolegend | 574704 | B255322 | 500 ng/ml |
| CX3CL1 | R&D Systems | 365-FR-025/CF | AF50521041 | 100 ng/ml |
| CCL2 | Biolegend | 571404 | B365559 | 50 ng/ml |
| CCL8 | Biolegend | 581604 | B370430 | 50, 100 ng/ml |
